# Distinct transcriptional programs define human IgG4^+^ memory B cells

**DOI:** 10.64898/2026.06.05.728165

**Authors:** Laurent M. Paardekooper, Jessica M. van Bokkum, Yvonne Fillié-Grijpma, Dana L.E. Vergoossen, Isabel Benner, Theresa Kissel, Susan L. Kloet, Martijn R. Tannemaat, Jan J. Verschuuren, Silvère M. van der Maarel, Maartje G. Huijbers

## Abstract

Human immunoglobulin G4 (IgG4) shapes both protective and pathogenic immunity, influencing conditions ranging from allergy and autoimmunity to cancer. However, subclass-specific targeting of memory B cells remains challenging due to limited knowledge of their phenotypic and functional heterogeneity. Here, we show that IgG4^+^ memory B cells are overrepresented in a niche of FCER2^+^BAFFR^+^ memory B cells, can be classified in several subtypes, have a distinct transcription factor profile and are the only memory B cell subset to express sterile *IGHE* transcripts. While IgG4^+^ B cells display a unique transcriptional profile, only BAFFR, IL5Rb and the B cell receptor were upregulated on protein level on IgG4^+^ cells. IgG4^+^ B cells have normal repertoire diversity, but a distinct germline V-gene usage, suggesting IgG4 responses are driven by specific antigens. By labeling autoreactive B cells, we confirmed that MuSK myasthenia gravis (an archetypal IgG4-mediated autoimmune disease) patients have a normal memory B cell profile and that autoreactive IgG4^+^ memory B cells are extremely rare. These results highlight the unique features of IgG4^+^ B cells, provide insight on potential subclass-specific therapeutic targets and point towards an antigen-driven IgG4 response within a largely non-autoreactive memory B cell pool.

## Introduction

Humoral immunity is a major branch of the adaptive immune system, protecting our body from a wide range of pathogens including viruses, bacteria, fungi and parasites. This broad range of threats requires an equally broad arsenal of antibody isotypes and subclasses. B cell responses typically start with IgM secretion, but as the response matures a subset of B cells undergoes class switch recombination to IgG1-4, IgA1-2 or IgE (*1*), coupled with affinity maturation and memory formation. This is regulated by a combination of T cell help in the germinal center, specific cytokine combinations and the nature and location of the antigen (*2*). Each subclass and isotype is characterized by a unique fragment crystallizable (Fc) domain, which is the main factor by which antibody effector functions are controlled. These mainly consist of complement deposition and Fc receptor-mediated activation of immune cells (*3–6*). In addition, a variety of Fc receptors is expressed across different immune ce **l** subsets, further finetuning their interactions with the humoral immune system by controlling antibody half-life, antigen presentation and immunomodulatory pathways (*7*, *8*).

The source of these diverse antibody responses lies within memory B cells (Bmems), which are a heterogenous and dynamic population of cells expressing surface bound antibodies called the B cell receptor (BCR). Activation of BCR signaling by a cognate antigen can drive memory B cells to class switching (*9*). As a consequence, B cells serve a range of effector functions depending on the isotype and subclass of the BCR, environment (germinal center or extrafollicular) and stimulation by T helper cells or follicular dendritic cells (*10*). Major Bmem effector functions include antibody secretion (by differentiation into short- or long-lived plasmablasts), antigen presentation to T cells or secretion of (regulatory) cytokines (*10*). This can be observed at the single cell level in both transcriptomic and proteomic profiling studies (*11*, *12*). Given the broad range of immune responses required to combat specific pathogens, memory B cells of each isotype and subclass are likely to all have a distinct phenotype.

While most immunoglobulins have potent inflammatory capacity, IgG4 is considered an anti-inflammatory antibody (*13*). This is due to its overall limited ability to interact with Fc receptors and low affinity for complement (*5*, *14*). IgG4 furthermore undergoes a stochastic process called Fab-arm exchange, in which it continuously exchanges half-antibody molecules with other IgG4 to become functionally bispecific, reducing antigen-specific avidity (*15*, *16*). We therefore hypothesize that class-switching to IgG4 is regulated via distinct pathways and possibly more strictly controlled given the potential risk of tolerogenic responses to harmful pathogens. Recently, IgG4, in spite of its anti-inflammatory nature, has been recognized to play a central (potentially) adverse role in a group of 29 autoimmune diseases (*17*), specific cancers (*18*), certain anti-drug antibody responses (*19*) and in mRNA vaccination (*20*, *21*). In contrast, IgG4 responses may alleviate allergic responses (*22*, *23*). Stimulating or dampening (antigen-specific) IgG4 responses therefore holds therapeutic promise.

To improve our understanding of the transcriptional and functional diversity within isotype-specific memory B cells and in particular that of IgG4 memory B cells, we generated unbiased single-ce **l** transcriptional profiles and V(D)J repertoire libraries by enriching for each isotype and subclass of memory B cells to build an atlas of isotype-specific marker genes in memory B cells. We also include similar data for MuSK MG patients, an archetypical neuromuscular autoimmune disease hallmarked by predominant IgG4 autoantibodies. We complemented these transcriptomic approaches with multi-color flow cytometry to deepen our understanding of the phenotype of these cells.

## Results

### Generation of a single-cell atlas of human isotype-specific memory B cells

To generate an atlas of isotype-specific memory B cells, we developed a sorting strategy to enrich for rare-isotype specific memory B cells using flow cytometry and performed single cell RNA sequencing (scRNA-seq) on sorted cells from three healthy donors and four MuSK MG patients (Fig. 1A, Sup. Table 1). Live Bmems were sorted from peripheral blood mononuclear cells (PBMCs) by staining for B cell markers (CD19, CD20) and selective depletion of IgM^+^ and IgG1^+^ Bmems combined with enrichment of IgG4^+^ Bmems Fig. 1B, Sup. Table 2). Cell sorting increased the IgG4^+^ cell count from 0.9% to 12% (Fig. 1C). 5’ Gene expression, CITE-seq-based MuSK-reactivity and full-length V(D)J (BCR-seq) sequencing libraries were constructed (*24*). After preprocessing and quality control (Sup. Fig. 1), the final integrated dataset includes 34.540 Bmems (15,164 cells from healthy controls and 19,376 cells from MuSK MG patients (Fig. 1D)). 8,388, 4,300 and 2,476 Bmems were sequenced from 3 healthy controls; 1,173, 4,423, 11,688 and 2,092 Bmems were sequenced from 4 MuSK MG patients, respectively. Unsupervised clustering of cells revealed 13 distinct clusters (Fig. 2A, Sup. Table 3) which were manually annotated using a selection of markers based on literature (Fig. 2B). In addition to classical subsets, several notable subsets were found, including a recently described subset of NKG7^+^GNLY^+^ cytotoxic Bmems (*25*) that also express genes encoding for granzyme B (*GZMB*) and perforin 1 (*PRF1*) and a small subset (58 cells) with high expression of interferon-activated genes such as *IFIT3*, *MX2*, and *IFI6*. We found a small but significant increase of this early cytotoxic subset for MuSK MG patients (Sup. Fig. 2). However, both the MuSK bait CITE-seq counts and the IgG4^+^ Bmem count within this subset were comparable between healthy donors and MuSK MG patients (Sup. Fig. 3), suggesting that these cells are unlikely to be direct drivers of autoreactivity in MuSK MG. Given the overlap between MuSK MG patients and healthy donors in cluster counts (Sup. Fig. 2) and differential gene expression (Sup. Fig. 4), we pooled the data for all following analyses.

**Figure 1.**
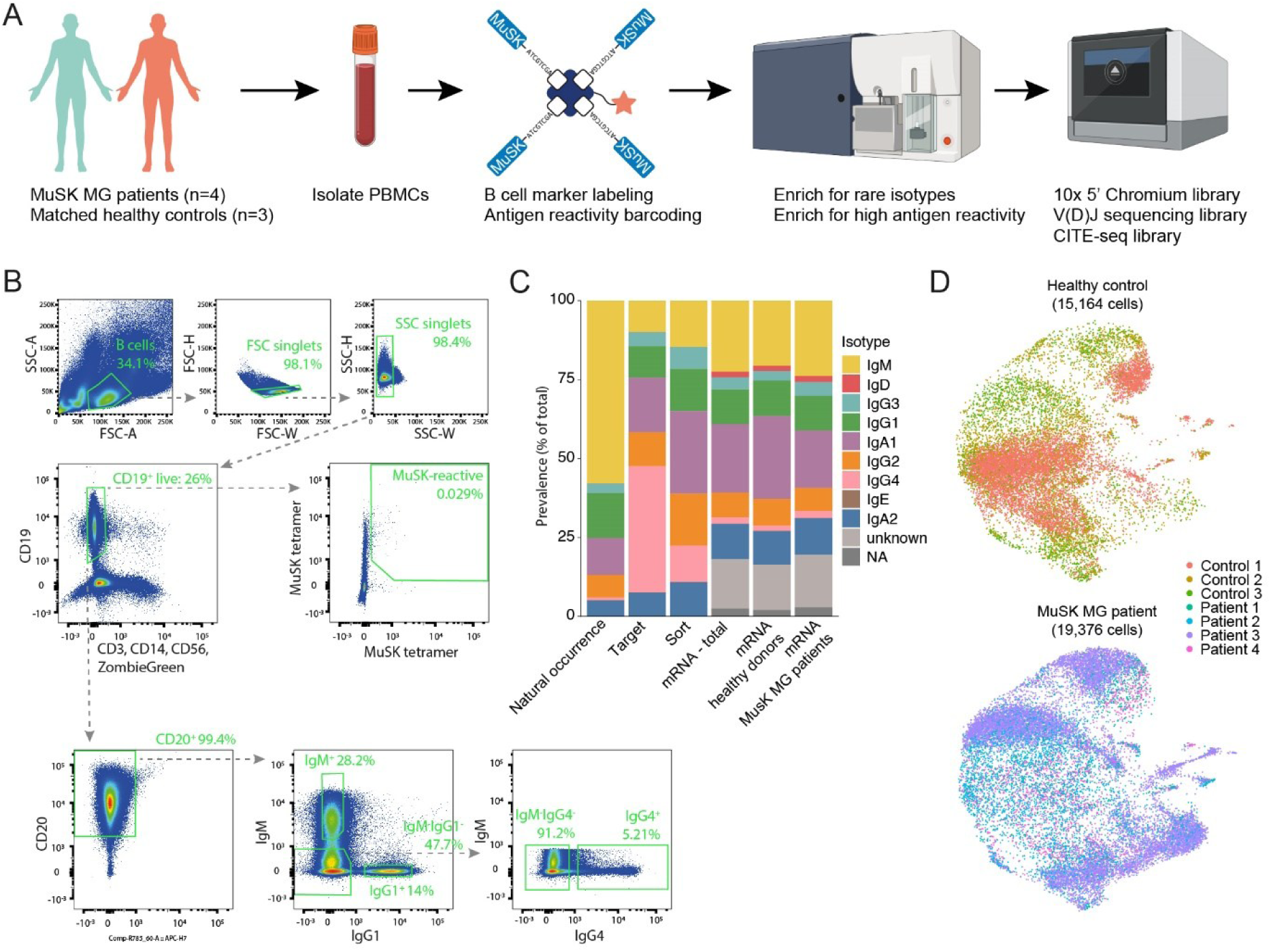
Generation of an isotype-specific single cell atlas of human memory B cells. **(A)** Schematic overview of the work flow for isolation, sorting and single cell transcriptomics experiments. **(B)** Representative dot plots of gating strategy to enrich for MuSK-reactive and IgG4^+^ memory B cells while depleting IgM^+^ and IgG1^+^ memory B cells. **(C)** Isotype distribution before and after IgM/IgG1 depletion and IgG4 enrichment. **(D)** Contributions of each donor to total UMAP projection.

**Figure 2.**
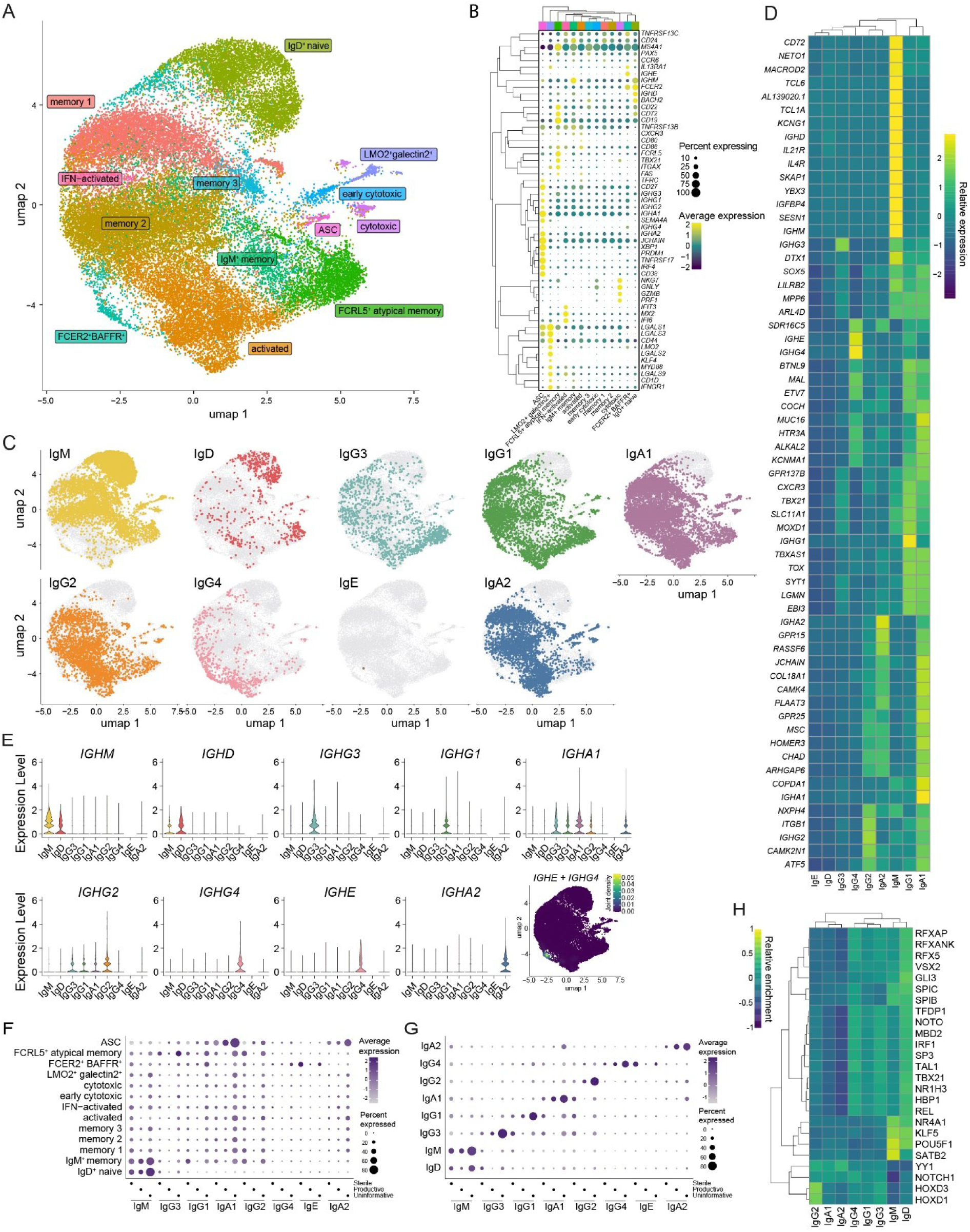
An isotype-specific single cell atlas reveals IgA^+^ and IgG4^+^ memory B cells have the most unique transcriptional profiles. **(A)** UMAP projection of complete dataset, clusters colored by B cell subset annotation. **(B)** Dot plot of marker gene expression shown for each annotated cluster shown in the UMAP (A). **(C)** Distribution of isotypes as determined by the V(D)J sequencing library over the combined UMAP plot shown in B. **(D)** Heatmap of the scaled relative expression of the top 15 differentially expressed genes per isotype. **(E)** Violin plots of normalized *IGH* gene expression per isotype as determined by the V(D)J sequencing library (x-axis). Kernel density estimation projection on UMAP of cells expressing both *IGHE* and *IGHG4*. **(F)** Expression of sterile, productive and uninformative heavy chain transcripts per Bmem isotype as determined by the 10X BCR-seq library. **(G).** Expression of sterile, productive and uninformative heavy chain transcripts per cell type as annotated in (B). **(H)** Heatmap of the scaled and relative inferred transcription factor activity per isotype as determined by the V(D)J sequencing library.

### IgA^+^ and IgG4^+^ Bmems are the most transcriptionally unique of all isotype-switched memory B cells

Next, we investigated representation of each immunoglobulin isotype and subclass as annotated by BCR-seq across the different clusters (Fig. 2A). As expected, IgHM^+^IgHD^+^ naive B cells cluster separately from the other Bmem clusters and express key markers of naive B cells (*IGHD, IGHM, BACH2, CD22, FCER2*) (*26–30*). IgG1-3^+^ and IgA1-2^+^ Bmems are found in all other clusters, while IgG4^+^ Bmems are strongly enriched in the memory 1, memory 2, FCER2^+^BAFFR^+^ and activated clusters, indicating IgG4^+^ Bmems have a distinct phenotype. Notably, only 1 IgE^+^ cell was recovered, which is expected given their relative rarity in blood (*31*) and that an IgE enrichment was not purused (Sup. Table 2). We investigated the top 10 upregulated genes for each isotype as annotated by BCR-seq and found the *IGH* heavy chain (HC) genes to be the main markers for each isotype (Fig. 2D). Furthermore, IgM^+^ Bmems show the most distinct gene expression profile with many uniquely expressed marker genes. IgG1^+^ Bmems share many marker genes with IgA1^+^ Bmems and show relatively high upregulation of *CXCR3*, *TBX21*, *SLC11A1* and *MOXD1*. IgG2^+^ Bmems highly express *NXXPH4* and *ITGB1*; although these genes were also (less strongly) upregulated in IgA1^+^ Bmems. The modest relative expression signature of IgG3^+^ Bmems overlaps with IgG1^+^ and IgA1^+^ Bmems. IgA1^+^ Bmems show a clear unique gene expression profile with high expression of, among others, *COPDA1*, *MUC16* and *JCHAIN* alongside the overlap with IgG1^+^ and IgG3^+^ Bmems mentioned above. Marker genes for IgA2^+^ Bmems mostly overlap with IgA1^+^ Bmems (*GPR15*, *RASSF6*) and IgG4^+^ Bmems (*SDR16C5*). No clear relative expression signatures were detected for IgD^+^ Bmems. IgG4^+^ Bmems share several upregulated genes with IgA1^+^ and IgG1^+^ Bmems and furthermore uniquely express *IGHE*, which matches a previous report on allergy-related Bmems (*32*). *SDR16C5* is found to be most upregulated on IgG4^+^ Bmems.

The expression of *IGHE* in IgG4^+^ Bmems prompted us to investigate the expression of multiple HC genes for each isotype (Fig. 2E). As expected, we observed co-expression of *IGHD* and *IGHM* in IgD^+^ and IgM^+^ Bmems. Additionally, there is low-level expression of *IGHA1* in IgG1-3^+^ and IgA2^+^ Bmems. Using kernel density estimation (*33*), cells co-expressing *IGHE* with *IGHG4* were found to localize within the cluster of FCER2^+^BAFFR^+^ Bmems. In conclusion, transcriptional reservoirs of other isotypes in Bmems are restricted to IgE and IgA being expressed in IgG4^+^ and IgG1-3^+^ Bmems, respectively. Following these observations, we set out to investigate whether class switching to IgG4 follows a distinct pattern. Expression of a sterile heavy chain gene (lacking VDJ gene segments) predisposes class switching events (*34*, *35*). Using the R package *sciCSR* (*36*), we extracted these reads from the raw data and investigated the expression of sterile and productive heavy chain genes. When stratified for cell type, we observed that FCER2^+^BAFFR^+^ Bmems highly express sterile *IGHE* transcripts, suggesting that they are poised to switch to IgE (Fig. 2F). When stratified for BCR-seq isotype call, we observe expression of sterile and uninformative (for example, only one read was mapped to C exonic regions) *IGHA1* and *IGHG2* transcripts in a subset of IgG3^+^, IgG1^+^, IgA1^+^ and IgG2^+^ cells. (Fig. 2G). Expression of sterile as well as productive *IGHM* by IgD^+^ cells is observed, as expected. The expression of multiple subclasses shown in Fig. 2E matches the sterile transcript expression pattern in Fig. 2G. Expression of sterile *IGHG4* transcripts is not increased in any of the subsets we identified, suggesting IgG4^+^ Bmems do not originate from a distinct Bmem subset or isotype subclass. Pseudotime reconstruction using *Monocle3* (*37*) did not reveal a distinct trajectory towards the FCER2^+^BAFFR^+^ or activated Bmem subsets where IgG4 is overrepresented, suggesting that IgG4^+^ Bmems can originate from a variety of subsets (Sup. Fig. 5).

To investigate isotype-specific differences in overarching regulatory profiles, we inferred transcription factor activity for each isotype from our single cell gene expression data using the *decoupleR* and *OmniPathR* packages (*38*, *39*) (Fig. 2H). This revealed 3 major activity clusters grouping IgG2^+^, IgA1^+^ and IgA2^+^ Bmems; IgG4^+^, IgG1^+^ and IgG3^+^ Bmems or IgM^+^ and IgD^+^ Bmems together. The combined activity of transcription factors Notch 1, Spi-B and Spi-C is unique to IgG4^+^ Bmems and may drive the observed gene expression profile. In conclusion, IgG4^+^ Bmems have a unique transcriptional and regulatory profile and are the only Bmem subtype acting as a reservoir for IgE transcripts.

### IGHG4, IGHE, CD74, KLF2 and SDR16C5 are the main marker genes of IgG4^+^ Bmems

As the modulation of IgG4 antibody responses is of clinical interest, we set out to find unique markers and pathway activity of IgG4^+^ Bmems with potential as therapeutic target. To compare the phenotype of IgG4^+^ Bmems with other isotypes, we performed differential gene expression analysis (DEG). After correction for multiple testing, 264 genes with a log2 fold change >2 versus all other isotypes (Fig. 3A) and 227 genes with a log2 fold change >2 versus other class-switched Bmems (Fig. 3B) were found. We compared the overlap in genes with a log2 fold change >1 in both analyses. Only a small fraction (3%; 42 genes) of all identified DEGs are exclusive to the IgG4-isotype switched comparison (Fig. 3D), the majority (72%; 917 genes) is found in both comparisons (Fig. 3A-B).

**Figure 3.**
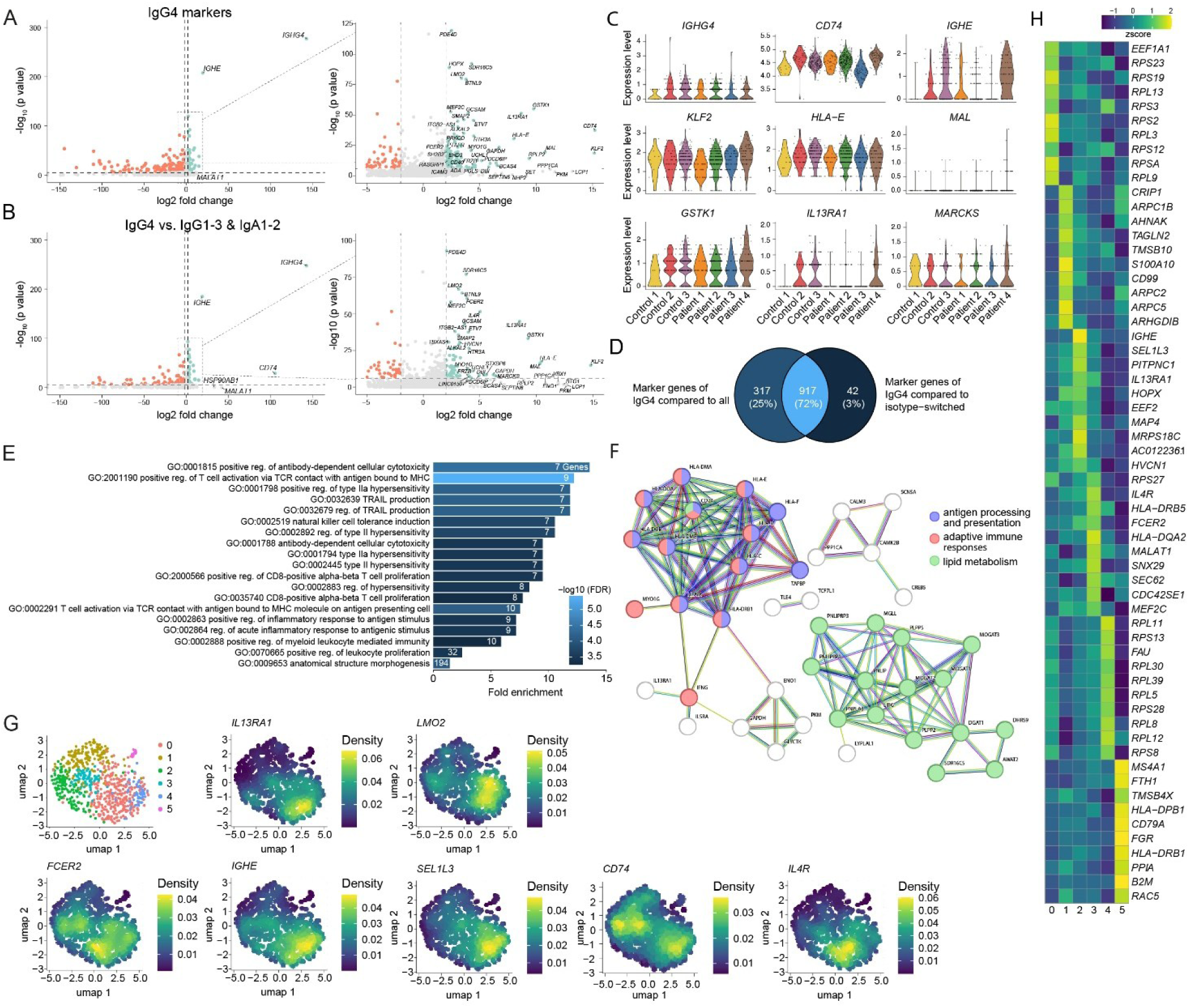
Unique expression profile of IgG4^+^ Bmems reveals a type II hypersensitivity-related subset. **(A-B)** Volcano plots showing differentially expressed genes of IgG4^+^ Bmems over all other Bmems (A) or of IgG4^+^ Bmems over isotype-switched Bmems (B). **(C)** Violin plot showing expression per donor of the major markers genes within IgG4^+^ cells found in A and B. **(D)** Overlap of marker genes found in (A) and (B). **(E)** Gene Ontology (GO) terms associated with the differentially expressed genes found in A. **(F)** STRING functional protein network of the top 100 differentially expressed genes found in A. **(G)** Reclustering of IgG4^+^ Bmems into a UMAP with kernel density estimations of the main differentially expressed genes found after reclustering. **(H)** Heatmap showing top 10 marker genes for each IgG4 subcluster identified in (G).

In both analyses, *IGHG4* and *IGHE* were the most significantly detected DEGs for IgG4^+^ Bmems by an order of magnitude. The gene with the highest log2 fold change not directly involved in BCR expression was *CD74*. This gene encodes for the class II MHC-associated invariant chain peptide (CLIP) and has several additional signaling functions in conjunction with CD44 and CXCR4 as a receptor for macrophage inhibiting factor (MIF) (*40–42*). Cytokine receptor-encoding genes *IL13RA1* and *IL4R*, both of which are associated with induction of IgG4 responses (*43*), were also significantly upregulated. *SDR16C5* encodes for Epidermal Retinol Dehydrogenase 2 (RDH-E2), an enzyme involved in retinoic acid biosynthesis (*44*), was also highly upregulated in IgG4^+^ Bmems. Retinoic acid is reported to increase IgG1^+^ B cell counts (*45*), induce switching to IgA (*46*, *47*) and upregulate CD1d expression to induce long-term memory responses (*48*). Its role in IgG4^+^ B cells is currently unknown. Furthermore, we found three transcription factors overexpressed in IgG4^+^ Bmems: *KLF2*, *LMO2* and *BTNL9*. These play a major role in B cell development (*49–51*) or have an immunomodulatory effect on T cells (*52–54*). Lastly, several factors affecting BCR signaling, B cell-T cell interactions and B cell survival, such as MARCKS and HOPX, where found upregulated in IgG4^+^ Bmems (*55–61*).

To investigate donor-specific variation in gene expression of the 9 most upregulated genes in Fig. 3A-B, they were individually plotted for each donor (Fig. 3C). All were found upregulated in both MuSK MG patient as well as healthy control B cells, and expression was confirmed for all donors although expression of *IL13RA1* and *IGHE* varied between samples (Fig. 3C). In conclusion, the DEG analysis identified a unique set of upregulated genes involved in BCR signaling, B cell survival and cytokines involved in IgG4 expression that is donor independent and IgG4^+^ Bmem specific. Next, to identify pathways uniquely active in IgG4+ Bmems, gene ontology (GO) analysis was performed on all genes upregulated in IgG4^+^ Bmems over Bmems of all other isotypes using ShinyGO 0.81 (*62*) (Fig. 3E). Selecting for biological processes with at least 5 upregulated genes involved, the major pathways upregulated in IgG4^+^ Bmems relate to antibody-dependent cellular cytotoxicity, T cell activation, antigen presentation to T cells and upregulation of type II hypersensitivity reactions. To investigate protein-protein association networks, we queried the STRING database for the top-100 upregulated genes in IgG4^+^ Bmems over Bmems of all other isotypes (*63*, *64*) (Fig. 3F). This analysis identified the main networks are formedby proteins involvedin antigen processing and presentation (GO: 0048002; blue in figure), adaptive immune responses (GO: 0002250; red in figure) and lipid metabolism (GO: 0006629; green in figure). Two additional, smaller networks were identified, but no GO terms were attached by the STRING database (white in figure). Combined, these results show IgG4^+^ Bmems have upregulated transcriptional programs for antigen presentation, T cell activation and the formation of T-B cell immunological synapses.

To investigate whether IgG4^+^ Bmem cells contain distinct subcluster populations, we performed unsupervised clustering, which yielded 6 unique clusters (Fig. 2F) and then used kernel density estimation (*33*) to identify unique marker genes (Fig. 3G). We assessed the top 10 marker genes expressed in each cluster identified in Fig. 3G (Fig. 3H). Only ribosomal genes were found in clusters 0 and 4. Cluster 5 only consists of 18 cells, limiting interpretation of the DEG analysis. Cluster 1 shows several differentially expressed cytoskeleton-related genes (several *ARP* genes, *TMSB10, ARHGDIB*). The dominant clusters 2 and 3 show high expression of *IGHE, IL13RA1, IL4R* and *FCER2*. This phenotype was described earlier to be closely related to allergic immunity (*32*). MuSK MG patient and healthy donor Bmems are equally represented in these clusters. IgG4^+^ B cells were also reported to contain a subset with proangiogenic properties (*65*), but we did not find any cells with this phenotype in our data. To conclude, IgG4^+^ Bmem subclusters can be classified largely in two groups that either display a type 2-polarized nature expressing *IGHE* transcripts and *FCER2* or those that do not have such a transcriptional profile, but rather appear at different conventional maturation or migration stages while expressing the IgG4 BCR.

### IgG4^+^ memory B cells have a distinct surface phenotype

To validate the markers found in our single cell RNA sequencing experiments on protein level, we carried out a flow cytometry screen on PBMCs obtained from seven healthy volunteers. Markers were selected from differentially expressed genes in Fig. 3A and Fig. 3B with known plasma membrane localization and high logfold change (Sup. Table 7). IgM^-^ live cells were gated on CD19^+^IgG4^+^, CD19^+^IgG4^-^ and CD19^-^IgG4^+^ subsets (Fig. 4A) and expression of the selected markers was assessed (Fig. 4B-C). Representative histograms from one donor are shown in Fig. 4B, with statistical analysis on seven donors shown in Fig. 4C. Notably, the CD19^+^IgG4^+^ subset shows higher expression of IgG4 (Fig. 4C) with a bipartite distribution (Fig. 4B). We identified IL5Rb (CD131) was specific to CD19^-^IgG4^+^ cells (Fig. 4B-C). IL5Rb, also known as common β chain, dimerizes with its co-receptor IL5Ra and is required for initiation of downstream signaling after stimulation with interleukin 5 (IL-5) (*66*, *67*). However, IL5Ra was not found to be upregulated on IgG4^+^ cells (Fig. 4B-C). IL13Ra1 (CD213a1) is significantly increased in the CD19^-^IgG4^+^ population and this trend is also visible in the CD19^+^IgG4^+^ population (Fig. 4B-C) and may indicate the presence of age-/autoimmune-associated B cells (*68*). In contrast to our transcriptional data, IL4R (CD124) and HLA-E surface levels were not detected in any of the subsets (Fig. 4C), precluding the formation of the IL4R-IL13Ra1 heterodimeric receptor complex in steady state. This complex is associated with induction of IgG4 responses (*69*, *70*). It is possible that protein expression of this complex is only induced after activation of IgG4^+^ Bmems, for example via the BCR, CD40 or BAFFR. IgE was only detected on the IgG4^+^ subsets (Fig. 4C), but the difference is not statistically significant (p = 0.0538) due to the large donor variation (Fig. 4C). FcεR2 (CD23), BAFFR (CD268) and CD74 were detected on both CD19^+^ subsets independent of IgG4 expression. BAFFR surface levels are significantly increased in IgG4^+^CD19^+^ Bmems, corroborating our transcriptomic findings (*TNFRSF13C*; Fig. 2A-C), but its presence is not exclusive to IgG4^+^ cells. GCSAM, CCR1 (CD191), MAL and TACI (CD267) were detected on all B cell subsets (Fig. 4C). In conclusion, the transcriptional markers observed on IgG4^+^ Bmems could only in part be confirmed on protein level for cell surface proteins. IL5Rb and the IgG4 BCR are the most differentially expressed cell surface protein markers for IgG4^+^ memory B cells.

**Figure 4.**
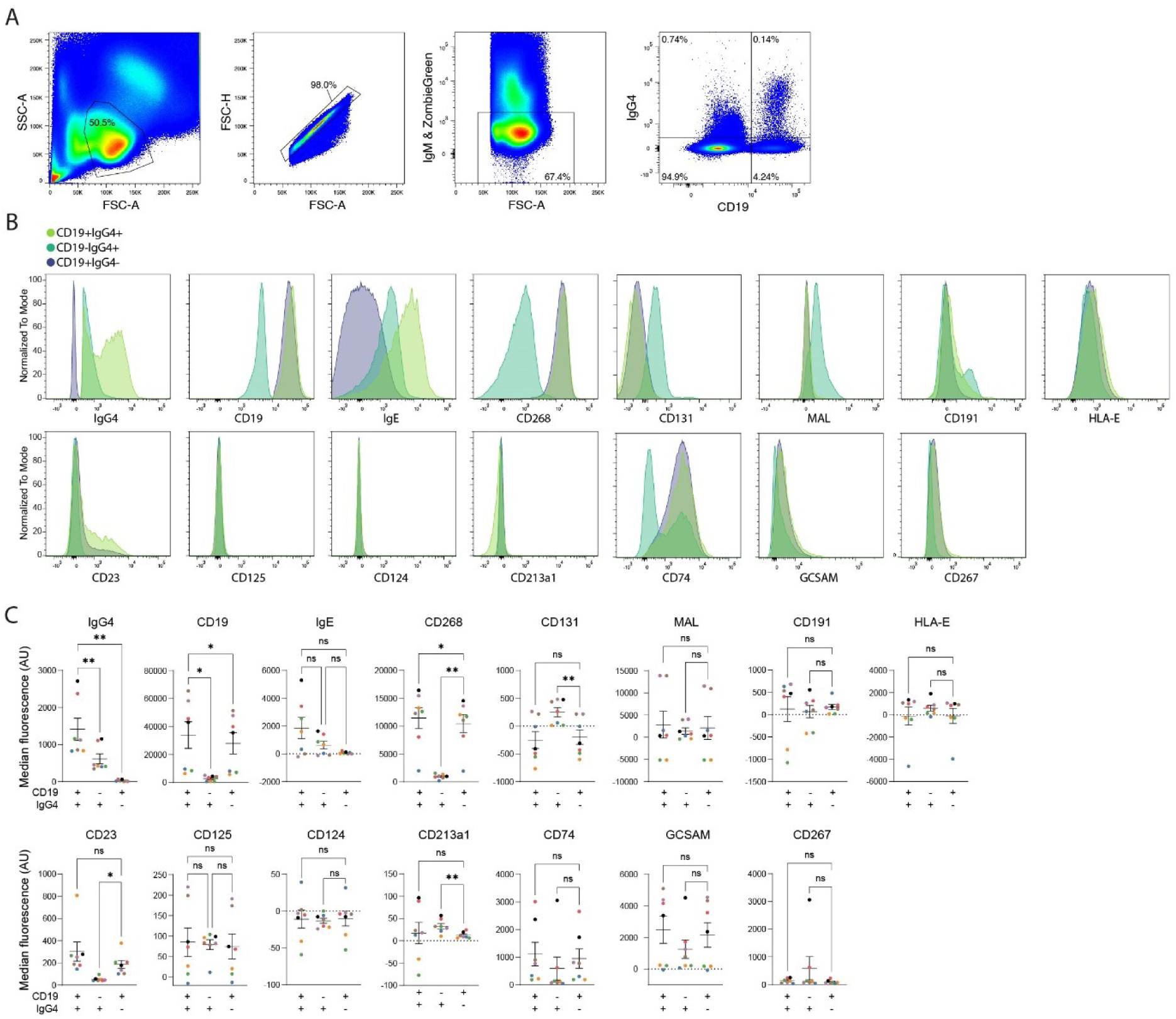
Surface marker validation of differentially expressed genes in IgG4+ B cells. **(A)** Representative histograms of flow cytometry gating before analysis of selected marker genes. **(B)** Representative flow histograms showing normalized expression of selected marker genes. **(C)** Flow cytometry validation of selected marker genes of 7 healthy donors (colors indicate paired data). *: P ≤ 0. 05, **: P ≤ 0. 005 (One-way ANOVA with Šidák correction for multiple comparisons).

### MuSK MG patients have a normal memory B cell compartment and autoreactive cells are extremely rare

To investigate whether patients with autoimmune disease predominated by IgG4 autoantibodies have an altered (IgG4^+^) Bmem compartment, we compared Bmem subset abundance and total Bmem transcriptomes between MuSK MG patients and matched healthy controls.

Differential gene expression analysis on MuSK MG patient versus healthy control Bmems revealed increased expression of mostly ribosomal genes and some immunological markers including *CD74, CST3, ACTB, IGHA1, IGHG1, GNLY* and several HLA-related genes (Sup. Fig. 4). *IGHG3, JCHAIN, MZB1, MALAT1* and several ribosomal genes are downregulated in MuSK MG patients (Sup. Fig. 4). Given the abundance of ribosomal genes, which are usually considered experimental background (*71*, *72*), combined with a lack of connecting pathways between these differentially expressed genes, we conclude there is no MuSK MG-specific gene expression pattern. Notably, there is also no difference in *IGHG4* expression between MuSK MG patients and healthy controls. Differential gene expression analysis on the IgG4^+^ Bmem subset revealed no immunologically relevant gene hits between MuSK MG patients and healthy controls (Sup. Fig. 4). Based on this data, we conclude that, overall, MuSK MG patients have a normal Bmem compartment. The pathogenic IgG4 autoantibody response is therefore likely to be antigen-driven.

To identify auto-reactive Bmems of MuSK MG patients in the scRNA-seq data, a multimodal, tetrameric, recombinant MuSK antigen bait (Fig. 5A) was included in the cell sorting step (Fig. 5A). This bait consists of the extracellular domains of MuSK (residues 1-486) fused to a C-terminal sortase motif (LPETGG) and 6His tag. Using recombinant *S. aureus* Sortase A pentamutant (*73*, *74*), we coupled this recombinant antigen to a 3’ biotinylated, CITE-seq compatible DNA barcode oligo (*24*) and subsequently tetramerized it using fluorescently-labeled streptavidin (Sup. Fig. 8). A negative bait tetramer was constructed in similar fashion by coupling a naked DNA barcode oligo to streptavidin with the same fluorescent labels (Fig. 6A). Both absolute and relative double-positive sort events were increased ± twofold in patient samples (representative plot shown in Fig. 5B). To validate these baits, a MuSK monoclonal antibody sequence (previously isolated from a patient (*75*)) was introduced as BCR in a MDL^-/-^AID^-/-^ RAMOS cell line (*76*, *77*) and validated with flow cytometry (Sup. Fig. 9). However, in our transcriptomics dataset, MuSK bait barcode counts of MuSK patient-derived Bmems were comparable to healthy controls despite correction for negative bait barcode counts (Fig. 5C). Cells with a MuSK barcode count ≥50 localize mostly in the IgD^+^, FCER2^+^ BAFFR^+^, memory 1 and activated clusters of MuSK MG patients and mostly in the IgD^+^, memory 2 and memory 3 clusters of healthy donors. We produced the BCR sequences of a selection of cells as recombinant antibodies to confirm their reactivity in a MuSK ELISA (*78*). Cells were selected based on MuSK barcode count, position in UMAP (Fig 2A, Fig. 5D), isotype and position in phylogenic tree (reconstructed using the Dowser package of the Immcantation suite (*79*, *80*)). The previously characterized 11-3F6 clone was used as a positive control (*75*). We confirmed 1 patient-derived IgG4 clone to be MuSK reactive (037-1-G4-1) (Fig. 5E). Phylogenetic trees were reconstructed from our BCR-seq data, from which a second set of clones was selected for screening based on their likeness to the 037-1-G4-1 clone previously confirmed as MuSK-reactive. However, none of the clones from this set were found to be MuSK-reactive (Fig. 5E). MuSK-reactive clones from MuSK MG patients are known to be extremely rare (order of 1-10/1·10^8^ PBMCs) (*75*, *81*, *82*). In total we tested 97 clones from 4 different patients, but beyond the first positive clone no other autoreactive B cells could be detected limiting further differential gene expression analysis between autoreactive cells and other Bmems.

**Figure 5.**
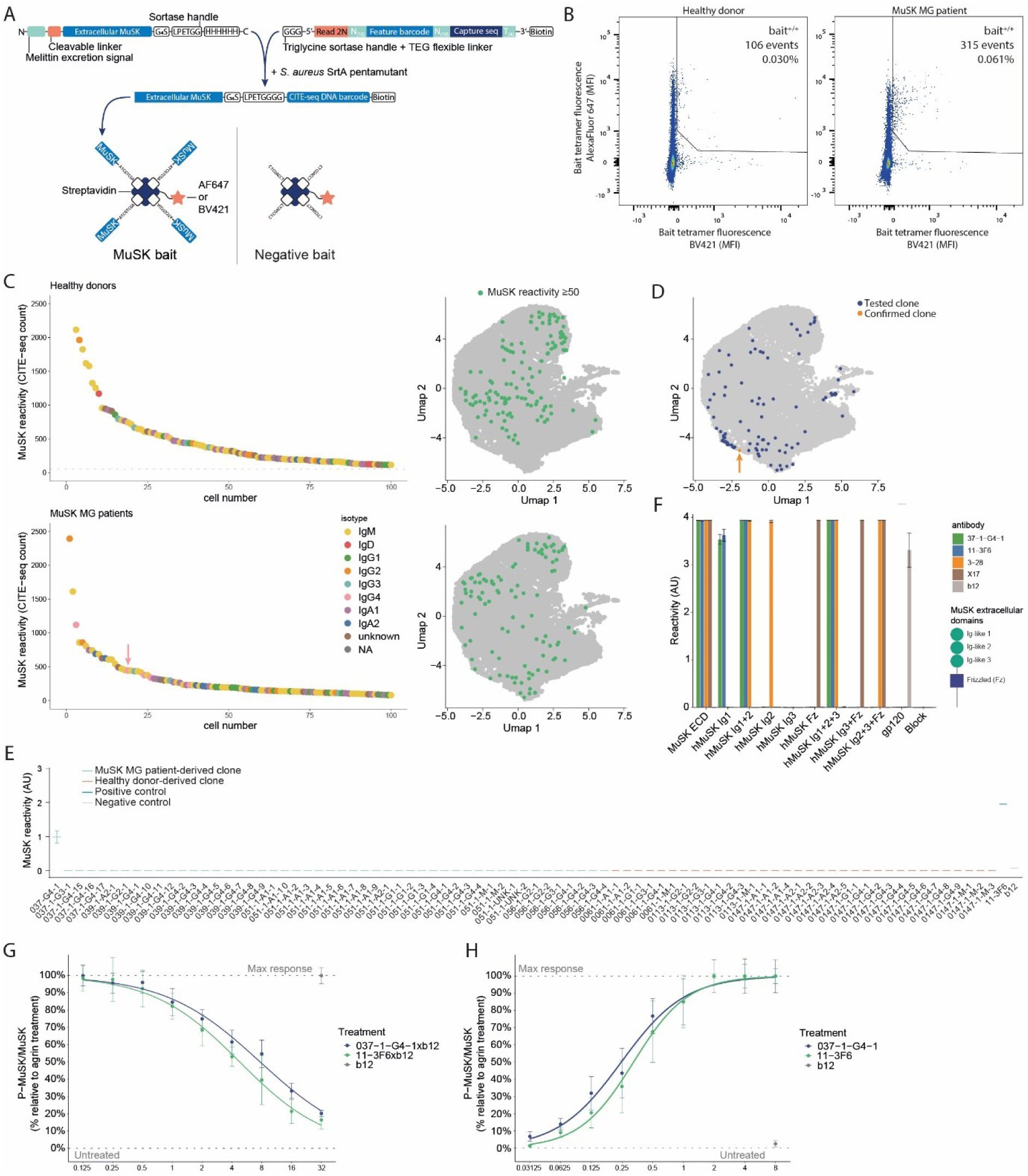
MuSK-reactive memory B cells are extremely rare. **(A)** Schematic overview of the tetrameric antigen baits used to identify MuSK-reactive cells in cell sorting and single cell RNA sequencing experiments. **(B)** Representative cell sorting dot plots of tetrameric antigen baits (MuSK bait and negative combined). Total event count and percentage of parent are displayed within the gate. **(C)** MuSK bait event counts in single cell RNA sequencing data after subtraction of negative bait counts. An arbitrary cutoff is placed at 50 events (dotted) line. All cells with ≥50 events are displayed on the UMAP also shown in Fig. 1B. Arrow indicates clone 037-1-G4-1. **(D)** UMAP position of cells selected for validation of BCR reactivity to MuSK. Orange dot indicates clone 037-1-G4-1. **(E)** MuSK ELISA of recombinant antibodies generated from selected BCR sequences. Error bars represent mean ± SEM. **(F)** MuSK epitope mapping ELISSA of MuSK-reactive clone 037-1-G4-1 identified in E. Clones 11-3F6, 3-28 and X17 All antibodies were tested at 1 μg/ml. Error bars represent mean ± SEM. **(G)** Inhibition of agrin-induced MuSK phosphorylation in C2C12 myotubes treated with monovalent anti-MuSK antibodies (n = 3). All data points were normalized to no antibody (100%) and untreated (0%) conditions. All antibody treatments are in the presence of 1 nM agrin. Error bars indicate mean ± SEM. **(H)** Induction of MuSK phosphorylation in C2C12 myotubes treated with bivalent anti-MuSK antibodies (037-1-G4-1: n = 3, 11-3F6: n = 2). All data points were normalized to 1 nM agrin control (100%) and untreated (0%) conditions. All antibody treatments are in the absence of agrin. Error bars indicate mean ± SEM.

**Figure 6.**
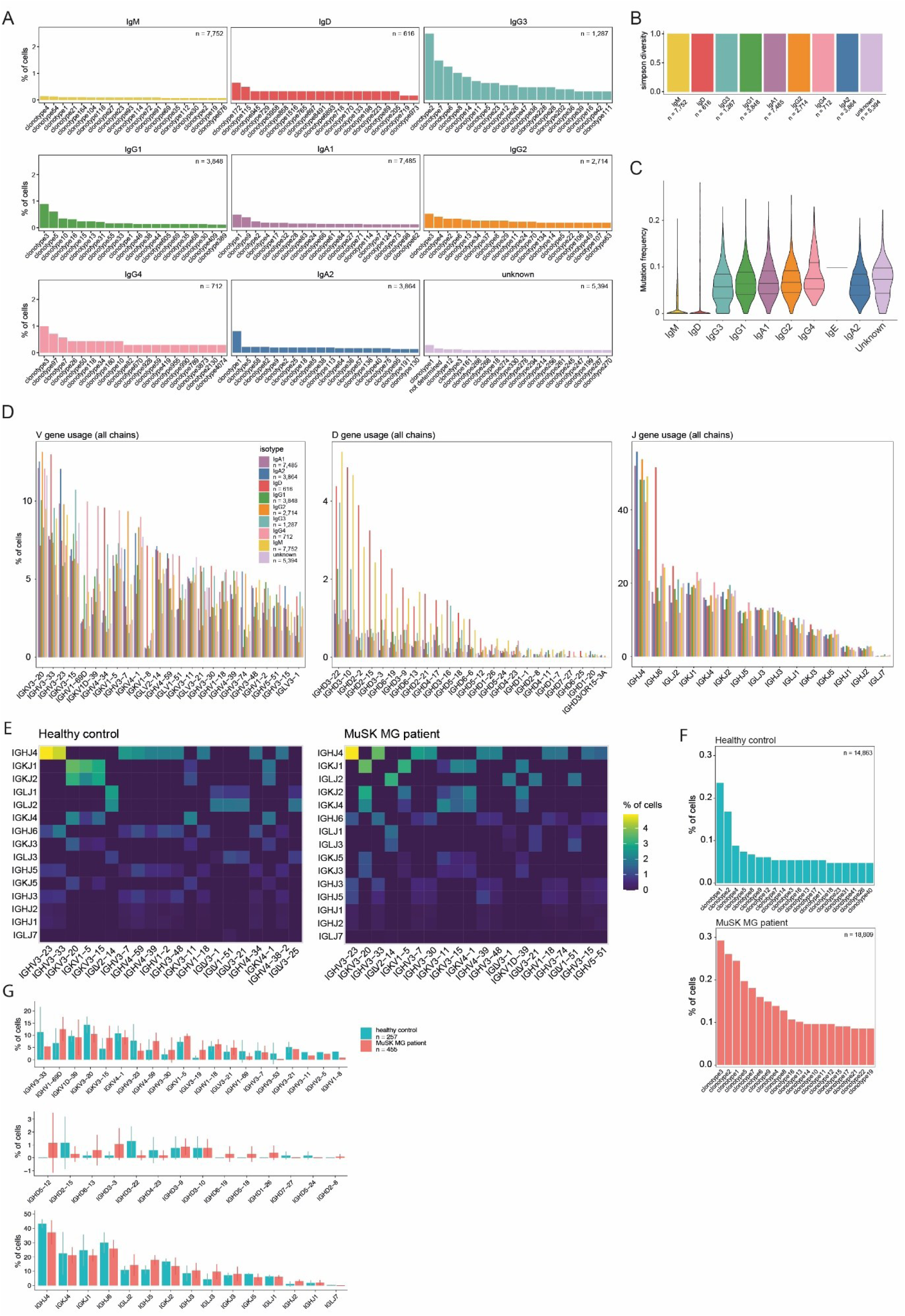
Repertoire analysis reveals comparable diversity between subclasses. **(A**) Isotype-specific clonotype analysis **(B)** Simpson diversity index calculated over all clones within each isotype. IgE was excluded due to lack of clones found **(C)** Somatic hypermutation (SHM) frequency for clones of each isotype calculated using the SHazaM package of the Immcantation suite. **(D)** V, D and J gene usage across heavy and light chains combined for each isotype. **(E)** Abundance of combined V-J gene usage (heavy and light chains combined) in healthy control and MuSK MG patient Bmems. **(F)** Clonotype analysis in healthy controls and MuSK MG patients. **(G)** Comparison of V, D and J gene usage in IgG4^+^ Bmems between healthy donors and MuSK MG patients.

Clone 037-1-G4-1 originates from an IgG4^+^ Bmem in the FCER2^+^BAFFR^+^ cluster (Fig. 2A), uses IGHV1-69*20 - IGHD3-3*01 - IGHJ3*01 and IGKV3-20*01 - IGKJ4*01, has a MuSK-reactivity score of 444 barcode reads and has a predicted N-glycosylation site at position 119. Epitope mapping revealed this clone binds to the Ig-like 1 domain of MuSK (*83*) (Fig. 5F). The Ig-like 1 domain is the interacting surface for LRP4 and agrin; which are essential interactions for activation of the downstreamsignaling cascade of MuSK, resulting in neuromuscular synaptic stability required for effective neuromuscular communication and muscle contraction (*84–86*). Previous studies have confirmed the pathogenicity of several other functionally monovalent anti-MuSKantibodies targeting the Ig-like domain 1 of MuSK in mice (*78*, *82*, *83*). To further investigate the pathogenic potential of the 037-1-G4-1 clone, we investigated its capacity to inhibit the agrin-Lrp4-MuSK signaling cascade (*87–89*) *in vitro* in C2C12 myotube cultures (*83*) (Fig. 5G). 037-1-G4-1 was subjected to controlled Fab-arm exchange with the HIV anti-gp120 antibody b12 (*90*) to generate a bispecific, functionally monovalent, 037-1-G4-1xb12 IgG4 antibody (*83*, *91*, *92*). 037-1-G4-1xb12 inhibited agrin-induced MuSK phosphorylation in a dose-dependent manner (Fig. 5G). Incontrast, the bivalent 037-1-G4-1 IgG4 antibody had an agonistic effect on MuSK phosphorylation in the absence of agrin (Fig. 5H). This is similar to observations done with other anti-MuSK clones (*75*, *83*, *93*). Therefore, this clone is predicted to be pathogenic after Fab-arm exchange (*94*). In summary, a tetrameric recombinant antigen-based bait can be used to label autoreactive Bmems in cell sorting and single cell transcriptomics experiments. However, these MuSK-reactive Bmems are extremely rare in the circulation and it is therefore diff icult to study their phenotype.

### IgG4 BCR repertoires are equally diverse but show distinct V gene usage compared to other isotypes

Recently it was shown that serum IgG4 clonality is relatively similar to that of IgG1 (*95*). However, rare isotypes and subclasses are thus far underrepresented in repertoire diversity studies (*96*). To investigate this, single cell BCR sequences were analyzed using the *DJVDJ* R package (Sheridan, 2023) and the Immcantation suite (Gupta et al., 2015; Vander Heiden et al., 2014) to compare clonal diversity, somatic hypermutation levels and V-J gene usage for each isotype. To investigate clonal diversity within each isotype and subclass, sequences were first grouped in clonotypes using Change-O (*97*) (Fig. 6A). Although several expanded clonotypes were uniquely found within IgG4^+^ Bmems (for example clonotypes 97, 26, 50, 118, 34 and 180), others with similar expansion rates are also found in IgG2^+^ Bmems (clonotype 3), IgG3^+^ Bmems (clonotype 7) and IgG1^+^ Bmems (clonotypes 3 and 10). The Simpson’s Diversity Index of all BCR sequences within each isotype and subclass approaches 1 (infinite diversity) (Fig. 6B). This suggests that the IgG4 BCR repertoire is as diverse as other isotypes.

We then quantified the VDJ mutation frequency within each isotype and found the increase in mutational load to follow class-switching order, with IgG4 BCRs having the highest frequency of mutation (Fig. 6C). Next, we investigated the V, D and J gene usage across both heavy and light chains within each isotype and subclass (Fig. 6D). V gene usage was found to coincide the most with specific isotypes and subclasses. *IGKV1-8, IGHV3-33* and *IGHV4-34* are overrepresented in IgM/D and IgG3 BCRs, while *IGKV3-20* is overrepresented in most class-switched BCRs. The overrepresentation of *IGHV3-33* and *IGHV4* family genes in naive B cells has been observed before (*98*)*. IGHV3-74* is more abundantly found in IgA1/2 and IgG2 BCRs. IgG4 BCRs are 2-10 times more likely to use *IGHV1-69D*, *IGKV1D-39* or *IGKV1-5*. These observations are not caused by V-gene bias in a single donor (Sup. Fig. 7).

Many D genes are overrepresented in IgM, IgD and IgG3 BCRs by a factor 2-4. However, this is likely caused by detection bias, as the D gene contains the majority of the highly variable CDR3 region (Rosner et al., 2001). More mature, class-switched BCRs accumulate more mutations in the CDR3 region, rendering it difficult to correctly call with computational germline reversion. J gene usage is comparable across all BCRs, with the exception of *IGHJ6* being overrepresented in IgD BCRs. Conversely, *IGHJ4* is underrepresented in IgD BCRs. *IGHJ4* usage is known to be increased in memory cells compared to naive B cells (*98*).

Next, we compared combined V-J gene usage on both heavy and light chain BCRs between healthy donors and MuSK MG patients (Fig. 6E), revealing no major alterations in V-J combinations in MuSK MG patient-derived Bmems. Overall clonotype diversity appears to be slightly lower in MuSK MG patients (Fig. 6F), but this data does not indicate a strong monoclonal response in these patients. The individual V, D and J gene usage in IgG4^+^ Bmems is comparable between MuSK MG patients and healthy controls (Fig. 6G). In conclusion, Bmems are, to a certain extent, predestined to become IgG4^+^ based on their germline V-gene usage, but this is not related to developing MuSK MG.

## Discussion

IgG4 antibodies fulfill a uniquely protective role in the immune system due to their functional monovalency combined with relative immunological inertia. Nevertheless, IgG4 is associated with a range of diseases, including 29 AIDs that are specifically mediated by IgG4 (*17*). Due to their low abundance, IgG4^+^ Bmems remain an understudied subset. Using flow cytometry-based enrichment of rare isotypes in combination with single cell transcriptomics, we characterized IgG4^+^ Bmems of healthy donors and MuSK MG patients (an archetypical IgG4-AID) and compared their characteristics to other (isotype-switched) Bmems. In addition to gene expression analysis, we included both full length BCR sequencing to compare the IgG4 BCR repertoire to other isotypes and antigen-specific labeling to characterize autoreactive B cells in MuSK MG patients.

Single-cell transcriptomic profiling revealed that Bmems from MuSK MG patients are largely transcriptionally comparable to matched healthy controls, corroborating our previous flow cytometry observations on B cell subset development (*100*), while providing a higher-resolution view at the single-cell level. This indicates that the autoantibody response in MuSK MG is antigen-driven rather than the result of aberrant B cell development, fitting with earlier observations (*100*) and the known predisposition for certain HLA haplotypes in IgG4-AID (*101*, *102*). These observations justifiedpooling patient and control memory B cells for downstream analyses focused on IgG4⁺ populations.

scRNA-seq furthermore enabled us to resolve subset-specific distributions, revealing that IgG4⁺ Bmems are selectively enriched within FCER2⁺BAFFR⁺ and activated Bmem niches, whereas other isotype-switched Bmems are broadly and homogeneously distributed across subsets, highlighting a previously unappreciated compartmentalization unique to IgG4⁺ B cells.

We also investigated the unique transcriptional features of IgG4^+^ Bmems and found they are characterized by a distinct gene expression profile, with the major differentially expressed genesbeing *IGHG4, IGHE, CD74, IL13RA1, IL4RA, FCER2, SDR16C5* and *KLF2*. CD74 plays a diverse role in the immune system. CD74 is also the receptor for macrophage inhibitory factor (MIF) (*40*), inducing B cell survival (*103*). MIF was recently discovered to be a biomarker in AChR myasthenia gravis correlating to disease severity (*104*, *105*). However, the overexpression of CD74 could not be corroborated with flow cytometry experiments, suggesting CD74 may be post-transcriptionally regulated in IgG4^+^ Bmems. Cytokine receptor-encoding genes *IL13RA1* and *IL4R*, both of which are associated with induction of IgG4 responses (*43*), were also significantly upregulated. Activation of the IL-4 receptor induces B cell maturation, survival and generation of memory (*106*, *107*). Signaling through the IL-13R-alpha-1 was shown to induce age-associated B cells (ABCs; also known as double-negative B cells) (*68*), which are a major source of autoantibodies in systemic lupus erythematosus (SLE) and correlate with disease activity (*68*, *108*, *109*). However, the IL-4R-alpha and IL-13R-alpha-1 can also form a heterodimer complex that drives type II inflammatory responses via STAT6 activation (*70*) with FCER2 expression and class switching to IgG4 as a key hallmark (*110*, *111*). While inferred STAT6 activity was not found upregulated in IgG4^+^ Bmems in our data (Sup. Fig. 6), the observed overexpression of *FCER2* (Fig. 3B) does suggest this pathway is active IgG4^+^ Bmems. *SDR16C5* encodes for an enzyme involved in retinoic acid biosynthesis (*44*) and was recently reported to be uniquely upregulated by IgG4^+^ Bmems (*112*). Retinoic acid is reported to increase IgG1^+^ B cell counts (*45*), induce switching to IgA (*46*, *47*) and upregulate CD1d expression to induce long-term memory responses (*48*). It will be interesting to investigate the role of *SDR16C5* in IgG4^+^ B cells. The transcription factor KLF2 plays a major role in pre B cell development (*49*). While its role in Bmems is unclear, it is likely to induce quiescence and tissue retention (*113–115*) driven by FOXO1 (*116*). However, FOXO1 activity was not found upregulated in IgG4^+^ Bmems in our data (Sup. Fig. 6). Pathway analysis on all differentially expressed genes revealed IgG4^+^ Bmems are more prone to induction of type II hypersensitivity responses and T cell activation via MHC:TCR signaling (Fig. 3E-F). In flow cytometry analyses, we identified IL5Rb, BAFFR and the BCR to be the only proteins with increased abundance in IgG4^+^ Bmems (Fig. 4C). This finding does not overlap with the previously described stringent dependance of IgG4 switching on IL-4/IL-13 signaling (*43*). Overexpression of IL5Rb, BAFFR and the BCR within IgG4^+^ Bmems is dependent on CD19 expression, further underlining the presence of functionally distinct memory subsets within the IgG4 response.

Another unique feature of IgG4^+^ B cells is the distinct upregulation of transcription factors Notch 1, Spi-B and Spi-C. Spi-B and Spi-C belong to the E26-transformation-specific (ETS) family of transcription factors and have opposing effects on B cell development via the BACH2 transcription regulator protein (*117–119*). We observed slightly higher inferred Spi-C activity in comparison to Spi-B, suggesting that IgG4^+^ Bmems are more prone to differentiate towards plasmablasts (*117*). Spi-C was shown to facilitate IL-4-induced germline IgE transcription in conjunction with STAT6 (*120*), which is also observed here. Both Spi-C and Notch 1 activity are known to drive differentiation towards antibody-secreting cells (*119–121*). The aforementioned upregulation of *SDR16C5* further confirms the possibility that IgG4^+^ Bmems are more prone to differentiate towards plasmablasts than other Bmems (*48*).

We furthermore provide a high-resolution view on subclass-specific B cell antibody repertoire diversity. This analysis revealed IgG4^+^ Bmems show a distinct V gene usage, of in particular *IGHV1-69D*, *IGKV1D-39* and *IGKV1-5*. Somatic hypermutation analysis previously reported that class-switching to IgG4 likely occurs mostly directly from IgM, with a minority coming from IgG1 (*96*, *122*). To further investigate the origin of IgG4 memory responses, we mapped class-switch recombination dynamics by investigating expression of sterile *IGH* transcripts across Bmem subsets or isotypes and subclasses. Our study did not reveal a distinct class switching preference of any specific isotype or Bmem subset towards IgG4. However, IgG4^+^ Bmems uniquely express high levels of sterile *IGHE* transcripts, coinciding with expression of other genes related to type 2 allergic responses, such as *FCER2, IL4R* and *IL13RA1* (*32*, *123*). Previous studies have also shown that IgE memory is mostly found in IgG1^+^ and IgG4^+^ Bmems (*32*, *124*, *125*). Sterile transcripts of other subclasses were overrepresented in some cases, most notably sterile *IGHA1* and *IGHA2* transcripts in the antibody-secreting Bmem subset and *IGHA2* transcripts in IgA1^+^ Bmems. However, these observations are not as specific as the link between IgG4 and IgE, suggesting that this IgE reservoir is a unique feature of IgG4^+^ Bmems. Taken together, these observations suggest that induction of IgG4 memory is coupled to specific properties of the antigen in combination with certain permissive cytokines and that IgG4 memory does not originate from a specific Bmem subset or isotype/subclass.

Our study both encountered and highlighted several (technical) challenges. The detection of MuSK-reactive cells using the tetrameric-barcoded antigen bait had limited success. Autoreactive B cells are known to be extremely rare (*75*, *99*). We analyzed high-quality transcriptomes of 19,376 Bmems from 4 MuSK MG patients. We estimate based on this and other studies that an average adult has approximately 300 MuSK-reactive Bmems. Therefore, isolation of sufficient cells for transcriptomic analysis with droplet-based single cell RNA sequencing methods is challenging and may require alternative antigen-bait methods or the study of plasmablasts and (bone marrow) plasma cells in the future, using for example the antibody capture method by Asrat *et al.* (*126*). Furthermore, calling of exact isotype and subclass is conflicting between BCR-seq and 5’ gene expression data in a selection of cells and for 16% a subclass or isotype could not be determined at all. Importantly, our study highlights the limited translatability of transcriptional observations to actual differences on protein levelin immune cells. This begs for cautious interpretation of single cell transcriptomics studies lacking proteomic validation and argues for this to be a requirement in publishing these types of datasets.

In summary, with this study, we generateda phenotypic and gene regulatory atlas of subclass-specific (IgG4^+^) memory B cells in the context of both IgG4-mediated autoimmunity and in the healthy condition. This dataset forms a resource to further study subclass-specific transcriptional differences of Bmems which can be leveraged to identify subclass-specific B cell targets for therapeutic purposes. We particularly investigated characteristic features of IgG4^+^Bmems and found that they are more distinct than other subclasses. These transcriptional and functional differences probably enable tighter control of these cells which is warranted because of the tolerogenic nature of IgG4.

## Materials and methods

### Study population

Four Dutch MuSK MG patients and three age-matched healthy controls were included in this study. All patients gave written consent according to the Declaration of Helsinki, and the study was approved by the Leiden University Medical Center ethics committee (registered under protocol P17-011). Patients were included based on positive diagnosis of MuSK MG and positive serological test for MuSK antibodies (*127*). Patients treated with immunosuppressants at time of blood draw were excluded.

### PBMC isolation

Peripheral blood mononuclear cells (PBMCs) were isolated from fresh Na-Heparin patient blood by Ficoll density gradient and cryopreserved until use.

### RAMOS MuSK-specific B cell lines

#### Plasmids

The pKat and pMIG2a plasmids were constructed as previously described (*77*). The heavy and light chains of the MuSK reactive IgG1κ antibody clone 11-3F6 (*75*) were inserted into the pMIG2a plasmid using InFusion HD master mix (Takara Bio) using the primers show in Sup. Table 8. DNA was isolated from overnight minipreps and insertion was confirmed by Sanger sequencing using the primers pMIG2a_gag_fw, pMIG2a_CH3_rv, GFP_out_fw and M13_rv (Sup. Table 8).

### Cell lines

The MDL^-/-^AID^-/-^ (IGHM, IGHD, IGLC and activation- induced cytidin deaminase (AID) knockout) Ramos B cell line was generated by dr He, University of Freiburg. Ramos cells were cultured in RPMI1640 (Thermo Fischer Scientific) with 10% fetal calf serum and 100 U/ml pen-strep (Thermo Fisher Scientific). Ramos cells express *SLC7a1* and were retrovirally transduced as previously described (*76*, *77*). Briefly, pKat and pMIG2a plasmids were cotransfected into Phoenix-ECO (ATCC CRL-3212) cells using Lipofectamine 3000 (Thermo Fischer Scientific). Phoenix-ECO cells were maintained for 48 hours in DMEM (Thermo Fischer Scientific) with 10% fetal calf serum before collection of retroviral supernatant, which were directly used for transduction into MDL^-/-^AID^-/-^ Ramos cells.

GFP expressing MDL^-/-^AID^-/-^Ramos cells were confirmed to be MuSK reactive by flow cytometry (Sup. Fig. 8). Cells were incubated at room temperature for 30 minutes with 1 µg/ml recombinant monomeric His-tagged MuSK generated as described below. Subsequently, cells were stained with Goat anti-MuSK (AF562; 1:1000, R&D Labs), Mouse anti-His (27E8 #2366, 1:1000, Cell Signaling Technology) or anti-human IgG1-DL650 (MH161-1, 1:200, Novus Biologicals) for 30 minutes on ice. Anti-MuSK and anti-His stained samples were incubated with Alexa647 secondary antibodies raised in donkey (A-21447 & A-31571 1:1000, Thermo Fischer Scientific) for 30 minutes on ice. All incubation steps in 50 µl PBS with 2% BSA. Cells were washed 2 times with 150 µl cold PBS between incubation steps. Samples were analyzed on an LSR-II flow cytometer (BD Biosciences). GFP positive cells were enriched using a FACSAria 3 cell sorter (BD Biosciences) for further culturing.

### Recombinant MuSK tetramers

#### Plasmid

The extracellular domains of MuSK (aa 1-486) were cloned into the pCPF2.08 vector using *in vivo* assembly (IVA) cloning (*128*) with the MuSK_insert_fw, MuSK_insert_rv, pCPF_IVA_rv and pCPF_IVA_fw primers. The C-terminal His-tag was deleted using MuSK_HISdel_fw and MuSK_HISdel_rv and an N-terminal sortase motif, G4S linker and 6His-tag were inserted using MuSK_SortHis_fw and MuSK_SortHis_rv (all primer sequences in Sup. Table 8). A map of the final plasmid is shown in Sup. Fig. 10.

### Bacmid transposition and transfection of Sf9 cells

The final MuSK plasmid was transformed into DH10α *E. coli* for bacmid transposition. Colonies were grown on agar plates containing kanamycin (50 µg/mL), tetracyclin (10 µg/mL), gentamycin (7 µg/mL), X-gal (80 µg/mL) and IPTG (40 µg/mL) for 3 days before blue/white screening. Bacmids were isolated from minipreps and transposition was confirmed by DreamTAQ PCR using M13 primers, successful transposition yields a 3.9 kb band. 0.5·10^6^ Sf9 cells were transfected with the bacmid using CellFectin reagent (Thermo Fischer Scientific; 10 μl bacmid solution, 5 μl CellFectin in 200 μl SFM-II medium; incubated for 10 minutes at RT) for 5 hours at 27°C before replacing medium. Supernatant containing virus was harvested after 48 hours and stored at 4°C.

### Protein production

3·10^6^ Sf9 cells were infected with 6 ml virus supernatant and cultured for 72 hours in 1.5 l of SFM-II medium (Thermo Fischer Scientific). Culture supernatant was aspirated and filtered with a .22 μm filter before loading on an NGC FPLC setup (Bio-Rad) for protein purification using a HisTrap FF crude 5 ml column (GE) at a flow speed of 5 ml/min. The column was equilibrated at 20 mM Tris/HCl, 200 mM NaCl, pH8. After loading, the column was washed with 20 mM Tris/HCl, 200 mM NaCl, 20 mM imidazole, pH8. Protein was eluted in 20 mM Tris/HCl, 200 mM NaCl, 200 mM imidazole, pH8. Column was stripped with 20 mM Tris/HCl, 200 mM NaCl, 500 mM imidazole, pH8.

Presence of recombinant MuSK was confirmed using Western blot. Column flowthrough, wash and eluted fractions were run a 10% SDS-PAGE gel for 20 minutes at 60V, then 90 minutes at 115V before blotting on a PVDF membrane (Danaher) for 90 minutes at 115V on ice. Membrane was blocked for 1 hour in 2% BSA (IgG-free, protease-free; Jackson Labs lot nr 121122) in PBS and stained with Goat anti-MuSK (AF562; 1:1000, R&D Labs) and Mouse anti-His monoclonal (27E8 #2366; 1:1000, Cell Signaling Technology) antibodies in blocking buffer overnight. Membrane was washed in PBS 0.02% Tween-20 and incubated with Donkey-anti-Goat IRDye800CW conjugated (LI-Cor, Lincoln, NE, USA; 1:15,000) and Donkey-anti-Mouse IRDye680RD conjugated (LI-Cor, Lincoln, NE, USA; 1:15,000) antibodies for 1 hour at RT before imaging on an Odyssey CLx imaging system (LI-Cor, Lincoln, NE, USA).

Elution fractions 1-25 were concentrated and buffer exchanged to 20 mM HEPES, 150 mM NaCl, pH 7.5. Initial concentration to 5 ml and buffer exchange were performed with a Centriprep 10 kDa MWCO spin column (Millipore). Final concentration was performed with an Ultracel 10 kDa MWCO (Amicon) to 2.5 ml. All centrifugation steps at 4°C with cold buffer. Samples were aliquoted to 100 μl, flash frozen in liquid N_2_ and stored at -80°C.

### Sortagging and tetramerization

A biotinylated 10x compatible barcode oligo (Sup. Fig. 8) was covalently attached to recombinant MuSK monomers by 16 hour incubation at room temperature with sortase pentamutant (*129*) in sortase buffer (500 mM Tris-HCl (pH 7.5), 1.5 M NaCl, 100 mM CaCl_2_) and subsequently purified on Talon metal affinity resin (Takara Bio) as previously described (*73*, *130*). Barcode oligo-conjugated MuSK has lost the 6His tag and will no longer bind to the Talon resin. The supernatant was then concentrated on a Vivaspin 500 filtration device with a molecular weight cut-off of 50 kDa (Sartorius) to remove excess unconjugated barcode oligos. The filter retentate was aliquoted and stored at -20°C until further use. Barcoded MuSK-oligo conjugates were tetramerized by 3 stepwise incubations with streptavidin (Alexa Fluor 647 or Brilliant Violet 510 conjugates) (increasing MuSK:streptavidin molar ratio from 10:1 to 20:1 to 30:1) for 10 minutes per step in the dark at room temperature, followed by a final additional incubation of 30 minutes. A negative bait to filter out aspecific binding was created by tetramerizing an unconjugated oligo to the same fluorescently-labeled streptavidins and following the same steps. A 1:1:1:1 molar ratio mix of these 4 tetramers was used in the staining panel described below.

### Cell sorting

Frozen PBMCs were thawed, washed once with cold PBS and stained for viability using 50 μl ZombieGreen staining solution (1:800; Biolegend) for 15 minutes in the dark on ice. Without washing, cells were blocked for 30 minutes in 130 μl PBS containing 2% BSA (Gibco) on ice. Cells were washed twice, followed by incubation with the sort staining antibody cocktail (Sup. Table 2) for 30 minutes on ice before washing twice. All wash and incubation steps were carried out using ice cold PBS containing 2% BSA. Finally, cells were resuspended in 150 μl PBS containing 2% BSA and sorted on a FACSAria 3 cell sorter (BD Biosciences). Briefly, we enriched for IgG4 while simultaneously depleting IgM and IgG1, and additionally selected every cell double positive for MuSK tetramers (APC and BV510) regardless of isotype (Fig. 1B-C). Strategies to sort all subclasses and isotypes with individual stainings were unsuccessful due the significantly extended duration of the sorting process, resulting in an unacceptable decline in cell quality. Cells were sorted into ice cold PBS containing 0.04% BSA and immediately transferred to the Leiden Genome Technology Center (LGTC) for single cell RNA library prep using 10X 5’ V2 chemistry (10X Genomics).

### Flow cytometry

Cryopreserved PBMCs were thawed, washed 2x with PBS and stained with ZombieGreen (1:800 in PBS; BioLegend) for 15 minutes on ice before blocking in 1x BD Horizon Brillian Stain buffer (BD Biosciences) for 30 minutes. Cells were split over two staining panels (Sup. Table 3). After staining, cells were washed 2xc with PBS and analysed on a Fortessa X-20 (BD Biosciences).

### Transcriptomics - pre-processing

Raw read alignment, unique molecular identifier counting and barcode counting were performed using the standard Cell Ranger (10X Genomics, V6.0.1) pipeline. Reads were aligned to the reference human genome GRCh38-2020-A for 5’ gene expression and individual barcodes and UMIs were counted. BCR sequencing reads were aligned to vdj_GRCh38_alts_ensembl-5.0.0 and linked to gene expression count tables by UMI as initial BCR assignments. Final assignment of isotypes per cell was performed using IMGT HighV-QUEST.

Count tables were then imported into Seurat v5 (*131*). To prevent V(D)J genes from affecting downstream 5’ gene expression analysis, the following genes were removed from the data set: *IG[HKL][VDJ]*, *IGKC*, *IGLC[1-7]*, *AC233755.1*, *AC233755.2* (encoding IGHV4-38-2), *IGLL*. Cells were filtered on read count, percentage of ribosomal reads and percentage of mitochondrial reads (using the *miQC* package (*132*)). Following data normalization and integration, principal component analysis (PCA) was used to reduce dimensionality and cells were clustered using the uniform manifold approximation and projection (UMAP) algorithm (*133*). Clusters with high expression of non-B cell marker genes T cell (*CD3, CD4, CD8*), NK cell (*GNLY,* NKG7) or monocyte (*LYZ, S100A8, S100A9*) combined with low expression of known memory B cell marker genes (*CD19*, *MSA4A1, CD27*) and lacking a BCR sequence were removed from the data set. Batch correction was performed in Seurat using the CCA algorithm.

### Transcriptomics - differential gene expression analysis

Differential gene expression was assessed on single cell level in Seurat v5 with FindMarkers using the MAST algorithm (*134*) with batch as latent variable, unless otherwise specified. Results were plotted in volcano plots using the *EnhancedVolcano* and *scCustomize* packages (*135*, *136*). Density kernel estimations for genes in small subsets were plotted using *Nebulosa* (*33*).

### Transcriptomics - pathway & transcription factor analysis

Differentially expressed gene sets were imported into the STRING network analysis tool (*63*, *64*). Gene Ontology (GO) term enrichment was assessed using ShinyGO version 0.81 (*62*). The top 30 Biological Process GO terms were selected with redundant terms removed. Transcription factor activity was inferred using the *decoupleR* package (*38*).

### Transcriptomics - B cell receptor analysis

Mutation frequencies in the CDR region and VDJ gene usage of all BCR sequences were analyzed using the *DJVDJ* package (*137*) and the *ShaZaM* package of the Immcantation suite (*97*, *138–140*). Lineage trees were inferred using the *Alakazam, Dowser, SCOPer* and *ShaZaM* packages (*97*, *141–144*) and visualized using *ggtree* (*145*). Class-switch recombination dynamics were inferred by comparing VDJ sequences of sterile and productive transcripts using the *sciCSR* package (*36*).

### Antibody reconstruction

Antibodies were produced from selected B cell receptor sequences to confirm MuSK reactivity. Heavy and light chain amino acid sequences were ordered at Twist Biologics and BioIntron using pcDNA3.1(+)-equivalent IgG1, IgG4, Igκ and Igλ2 expression vectors. An antibody leader sequence (MDWTWSILFLVAAPTGAHS) was added to the N-terminus of the variable regions before codon optimization for *Homo sapiens*. Next, an EcoRI restriction site (GAATTC) and partial Kozak sequence (GCCACC) were added to the 5’ end of the antibody leader sequence. After transformation and midiprep, a pilot of 10 matching heavy and light chain plasmids were cotransfected in HEK293T FreeStyle cells and supernatants were tested for total IgG levels and MuSK reactivity using ELISAs. Further clones were directly produced at BioIntron with comparable methods.

### Epitope mapping ELISA

Antibodies were produced in IgG4-S228P backbones at BioIntron as described above, except for X17 which was produced as IgG1. Recombinant TwinStrep and FLAG double-tagged human MuSK epitopes were produced by BioIntron as described above (MuSK-ECD and gp120 only carried a TwinStrep tag). ELISA plates were coated with 3 ug/ml per well of epitope diluted in blocking buffer (2% casein, 0.05% tween-20 in PBS). After washing, specific binding was detected with an alkaline phosphatase-conjugated rabbit anti-human IgG (Jackson ImmunoResearch Europe 309-055-008; 1:1000 in blocking buffer) combined with 1 mg/ml PNP DEA substrate. Reaction was stopped with 3 M NaOH after 60 minutes and optical density was read at 405 nm. All steps were carried out for 60 minutes at room temperature on an orbital shaker (960 rpm).

### MuSK phosphorylation experiments

Antibodies were purified with protein A affinity chromatography as previously described (*75*). Functionally monovalent IgG4 MuSK antibodies were generated through controlled Fab-arm exchange with b12 hIgG4-S228P-F405L-R409K (*83*). C2C12 myoblast (ATCC C2C12 CRL-1772, lot 70058257) were maintained in DMEM (Thermo Fisher Scientific #31966-021) with 10% fetal bovine serum (FBS) (Sigma T7524). Myoblasts were differentiated into myotubes by seeding them at 1.40·10^4^ cells/cm^2^ in 24-well plates in 0.75 mL medium. Half of the medium volume was refreshed after 4-5 days. Seven days after seeding, C2C12 myotubes were treated with a 2-fold serial dilution of each antibody for 30 min at 37°C. For monovalent antibodies, 1 nM agrin (R&D systems 550-AG) was simultaneously added. After treatment, cells were washed once with 300 µL ice-cold PBS before incubation of 30 min (shaking at 4°C) with 100 µL of lysis buffer (50 mM Tris-HCl, 150 mM NaCl, 1mM EDTA, 1% NP-40, 0.25% deoxycholic acid, pH=7.4, supplemented with protease and phosphatase inhibitors (Merck Life science #4906837001, Roche #11836170001). Lysates were stored at -80°C until further use.

### MuSK phosphorylation ELISA

Phosphorylated and total MuSK were detected in the lysates with a binding ELISA based on previously described methods (*146*). Specifically, plates were coated overnight with MuSK-Fz domain antibody 30A11 (1 µg/mL in PBS, kind gift from argenx). Lysates were diluted 10x in 0.1% casein in PBS for the detection of total MuSK, used undiluted for phosphorylated MuSK, and tested in triplicate for each measurement. Total and phosphorylated MuSK were detected with respectively MuSK-Fz domain antibody X17 (1 µg/mL, BioIntron, in-house biotinylated), or a mix of biotinylated phosphotyrosine clone 4G10 (1 µg/mL, Merck Millipore Corp 16-103) and clone PY20 (1 µg/mL, Biolegend 309304), followed by streptavidin poly-HRP (Pierce PIER21140; 10 ng/mL for Total MuSK, 20 ng/mL for phosphorylated MuSK). Phosphorylated MuSK signals were first normalized to the total MuSK signal, followed by normalization to the response in agrin-treated (100%) and untreated (0%) myotubes per biological replicate. The normalized data was fitted to a four-parameter nonlinear regression model.

### Statistical analyses

Statistical analyses were performed in R using the MAST algorithm (*134*) for differential gene expression analysis called via EnhancedVolcano package (*135*). Flow cytometry data was analyzed using FlowJo v10. Statistical analyses on flow cytometry data were performed using GraphPad Prism 10.2.3. P-values and tests used are indicated in figure captions.

## Code and data availability

The code for this manuscript is available on Git: https://git.lumc.nl/lmpaardekooper/subclass-seq. The full transcriptomic dataset is being made available via the European Genome-Phenome Archive. In compliance with European privacy legislature, data access is available upon request.

## Acknowledgements

The authors gratefully acknowledge dr. Robbert Kim for his advice and support with generating recombinant MuSK protein.

The authors gratefully acknowledge dr. William F Jiemy for conducting the epitope mapping ELISA experiments in Figure 6.

The authors gratefully acknowledge dr. Ahmed Mahfouz, dr. Linda Slot, prof. dr. Ramon Arens and dr. Cees van Bergen for their advice and support with study design, data analysis and manuscript writing.

The authors gratefully acknowledge the Flow cytometry Core Facility (FCF) of Leiden University Medical Center (LUMC) in Leiden, the Netherlands (https://www.lumc.nl/research/facilities/fcf), coordinated by dr. K. Schepers and M. Hameetman, run by the FCF Operators E.F.E de Haas, J.P. Jansen, D.M. Lowie, S. van de Pas, and G.IJ. Reyneveld, directed by prof. dr. F.J.T. Staal) for technical support.

The authors gratefully acknowledge the Leiden Genome Technology Center (LGTC) of Leiden University Medical Center (LUMC) in Leiden, the Netherlands (https://www.lgtc.nl/), coordinated by dr. Susan Kloet, for technical support.

The authors gratefully acknowledge the LUMC Vrijwillige Donoren Service (LuVDS) and its volunteers for providing healthy donor material.

pet30b-5M SrtA was a gift from prof. dr. Hidde Ploegh (Addgene plasmid # 51140; http://n2t.net/addgene:51140;RRID:Addgene_51140).

MDL^-/-^AID^-/-^ RAMOS cell line was a kind gift from prof. dr. René Toes, dr. Theresa Kissel and prof. dr. Michael Reth.

GGG-6FAM dye was a kind gift from dr. Martijn Verdoes. Biorender was used in the generation of several figures.

## Author contributions

**L.M. Paardekooper:** methodology, software, formal analysis, investigation, writing – original draft, data curation, visualization

**J.M. van Bokkum:** software, formal analysis, data curation

**Y. Fillié-Grijpma:** investigation

**D.L.E. Vergoossen:** methodology, investigation

**I. Benner:** investigation

**T. Kissel:** resources

**S.L. Kloet:** methodology

**M.R. Tannemaat:** resources

**J.J. Verschuuren:** resources, supervision, writing – review & editing

**S.M. van der Maarel:** supervision, writing – review & editing

**M.G. Huijbers:** conceptualization, supervision, writing – review & editing, project administration, funding acquisition

## Funding

This project is financially supported by a Human Genetics Onderzoekscommissie (HGOC) NGS pilot grant 2021 (SG20210309).

MGH receives financial support from the LUMC Gisela Thier Fellowship 2021, ZONMW VENI 2019 (0915016181 0040) and the European Union (ERC, IgG4-START, 101163002). Views and opinions expressed are however those of the author(s) only and do not necessarily reflect those of the European Union or the European Research Council. Neither the European Union nor the granting authority can be held responsible for them.

The LUMC forms part of the European Reference Network for Rare Neuromuscular Diseases [ERN EURO-NMD] and the Netherlands Neuromuscular Disorders Center (NL-NMD).

## List of supplementary materials

Supplementary Table 1. Study population

Supplementary Table 2. Sort antibody panel

Supplementary Table 3. Cell counts per cluster for each donor

Supplementary Table 4. Cell counts per cluster for each group

Supplementary Table 5. Isotype counts per cluster for each donor

Supplementary Table 6. Isotype counts per cluster for each group

Supplementary Table 7. Flow cytometry antibody panels

Supplementary Table 8. Primers

Supplementary Figure 1. Single cell RNA seq quality control

Supplementary Figure 2. Relative abundance of MuSK MG patient and healthy control B cells

Supplementary Figure 3. MuSK reactivity scores and isotype distribution in the GNLY^+^NKG7^+^ Bmem subsets

Supplementary Figure 4. Differential gene expression of MuSK MG patient B cells over healthy controls

Supplementary Figure 5. Pseudotime inference using Monocle3 with IgD+ naive memory B cell cluster as starting node

Supplementary Figure 6. Inferred activity of transcription factors FOXO1 and STAT6 per isotype.

Supplementary Figure 7. Recombinant MuSK tetrameric antigen

Supplementary Figure 8. Validation of MuSK-reactive RAMOS cell line

Supplementary Figure 9. Plasmid map of recombinant extracellular sortaggable MuSK plasmid

**Supplementary Table 1.**
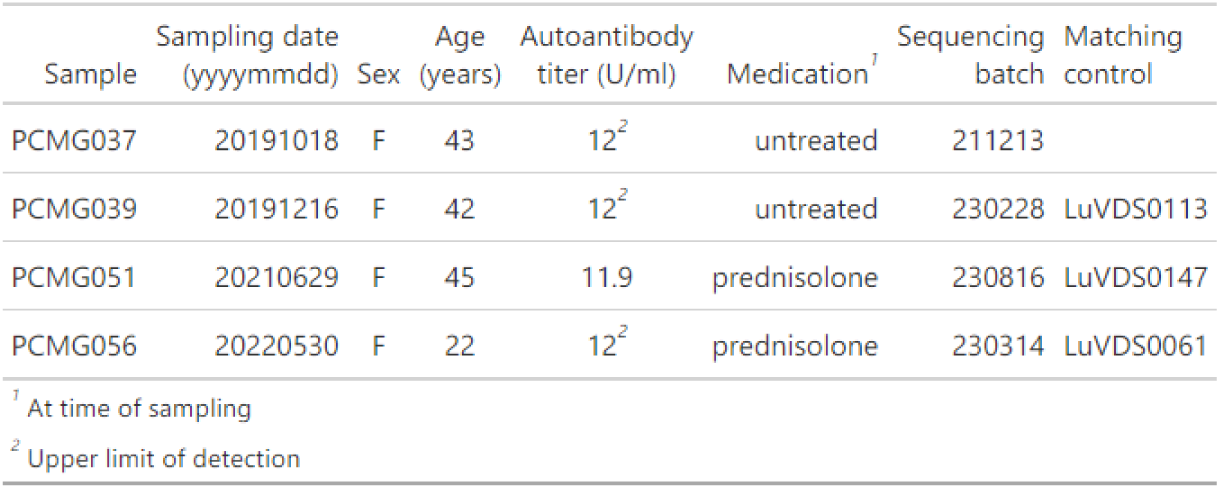
Study population.

| Sample | Sampling date<br>(yyyymmdd) | Sex | Age<br>(years) | Autoantibody<br>titer (U/ml) | Medication<sup>1</sup> | Sequencing<br>batch | Matching<br>control |
| --- | --- | --- | --- | --- | --- | --- | --- |
| PCMG037 | 20191018 | F | 43 | 12<sup>2</sup> | untreated | 211213 |  |
| PCMG039 | 20191216 | F | 42 | 12<sup>2</sup> | untreated | 230228 | LuVDS0113 |
| PCMG051 | 20210629 | F | 45 | 11.9 | prednisolone | 230816 | LuVDS0147 |
| PCMG056 | 20220530 | F | 22 | 12<sup>2</sup> | prednisolone | 230314 | LuVDS0061 |

<sup>1</sup> At time of sampling
<sup>2</sup> Upper limit of detection

**Supplementary Table 2.** Sort antibody panel.

| Marker | Clone | Dilution | Source | Cat. no |
| --- | --- | --- | --- | --- |
| CD19-BV605 | SJ25C1 | 1:50 | BioLegend | 363024 |
| CD20-PE-Cy7 | 2H7 | 1:100 | BioLegend | 302312 |
| IgM-AF700 | MHM-88 | 1:25 | BioLegend | 314538 |
| IgG1-PE<sup>1</sup> |  | 1:33 | Sanquin | M1325 |
| IgG4-DL594<sup>2</sup> | HP6098 | 1:33 | Sanquin | M1271 |
| Strep-AF647<sup>3</sup> | n.a. | 1:22 | BioLegend | 405237 |
| Strep-BV510<sup>3</sup> | n.a. | 1:22 | BioLegend | 405234 |
| CD3-FITC | UCHT1 | 1:100 | BD Biosciences | 561806 |
| CD14-FITC | M5E2 | 1:100 | BD Biosciences | 561712 |
| CD56-FITC | HCD56 | 1:100 | BioLegend | 318304 |
| ZombieGreen | n.a. | 1:800 | BioLegend | 423111 |

<sup>1</sup> Fluorophore directly conjugated using PE labeling reagent (Thermo Fisher Scientific)
<sup>2</sup> Fluorophore directly conjugated using NHS Ester labeling reagent (Thermo Fisher Scientific)
<sup>3</sup> Streptavidin bound to either recombinant barcoded MuSK antigen or negative bait oligo

**Supplementary Table 3.** Cell counts per cluster for each donor.

| UMAP cluster | Healthy donor |  |  |  |  |  | MuSK MG patient |  |  |  |  |  |  |  |
| --- | --- | --- | --- | --- | --- | --- | --- | --- | --- | --- | --- | --- | --- | --- |
|  | LuVDS0061 | LuVDS0061 (%) | LuVDS0113 | LuVDS0113 (%) | LuVDS0147 | LuVDS0147 (%) | PCMG037 | PCMG037 (%) | PCMG039 | PCMG039 (%) | PCMG051 | PCMG051 (%) | PCMG056 | PCMG056 (%) |
| memory 1 | 75 | 0.89 | 761 | 17.70 | 670 | 27.06 | 220 | 18.76 | 1131 | 25.57 | 3359 | 28.74 | 178 | 8.51 |
| activated | 1389 | 16.56 | 926 | 21.53 | 611 | 24.68 | 175 | 14.92 | 897 | 20.28 | 2435 | 20.83 | 505 | 24.14 |
| memory 2 | 4804 | 57.27 | 1439 | 33.47 | 482 | 19.47 | 248 | 21.14 | 1303 | 29.46 | 120 | 1.03 | 798 | 38.15 |
| IgHD+ naive | 1187 | 14.15 | 524 | 12.19 | 346 | 13.97 | 276 | 23.53 | 254 | 5.74 | 2162 | 18.50 | 288 | 13.77 |
| FCRL5+ atypical memory | 261 | 3.11 | 77 | 1.79 | 74 | 2.99 | 34 | 2.90 | 251 | 5.67 | 1812 | 15.50 | 45 | 2.15 |
| IgM+ memory | 357 | 4.26 | 222 | 5.16 | 108 | 4.36 | 73 | 6.22 | 329 | 7.44 | 571 | 4.89 | 84 | 4.02 |
| FCER2+ BAFFR+ | 124 | 1.48 | 226 | 5.26 | 128 | 5.17 | 53 | 4.52 | 128 | 2.89 | 264 | 2.26 | 93 | 4.45 |
| memory 3 | 33 | 0.39 | 32 | 0.74 | 17 | 0.69 | 27 | 2.30 | 37 | 0.84 | 336 | 2.87 | 38 | 1.82 |
| early cytotoxic | 16 | 0.19 | 20 | 0.47 | 12 | 0.48 | 32 | 2.73 | 26 | 0.59 | 361 | 3.09 | 15 | 0.72 |
| LMO2+ galectin2+ | 45 | 0.54 | 2 | 0.05 | 7 | 0.28 | 2 | 0.17 | 11 | 0.25 | 163 | 1.39 | 16 | 0.76 |
| cytotoxic | 51 | 0.61 | 47 | 1.09 | 4 | 0.16 | 18 | 1.53 | 35 | 0.79 | 62 | 0.53 | 21 | 1.00 |
| ASC | 38 | 0.45 | 15 | 0.35 | 5 | 0.20 | 3 | 0.26 | 10 | 0.23 | 41 | 0.35 | 7 | 0.33 |
| IFN-activated | 8 | 0.10 | 9 | 0.21 | 12 | 0.48 | 12 | 1.02 | 11 | 0.25 | 2 | 0.02 | 4 | 0.19 |

**Supplementary Table 4.** Cell counts per cluster for each group.

| UMAP Cluster | Healthy donor |  | MuSK MG patient |  |
| --- | --- | --- | --- | --- |
|  | Count | Percentage | Count | Percentage |
| memory 1 | 1506 | 9.93 | 4888 | 25.23 |
| activated | 2926 | 19.30 | 4012 | 20.71 |
| memory 2 | 6725 | 44.35 | 2469 | 12.74 |
| IgHD+ naive | 2057 | 13.57 | 2980 | 15.38 |
| FCRL5+ atypical memory | 412 | 2.72 | 2142 | 11.05 |
| IgM+ memory | 687 | 4.53 | 1057 | 5.46 |
| FCER2+ BAFFR+ | 478 | 3.15 | 538 | 2.78 |
| memory 3 | 82 | 0.54 | 438 | 2.26 |
| early cytotoxic | 48 | 0.32 | 434 | 2.24 |
| LMO2+ galectin2+ | 54 | 0.36 | 192 | 0.99 |
| cytotoxic | 102 | 0.67 | 136 | 0.70 |
| ASC | 58 | 0.38 | 61 | 0.31 |
| IFN-activated | 29 | 0.19 | 29 | 0.15 |

**Supplementary Table 5.**
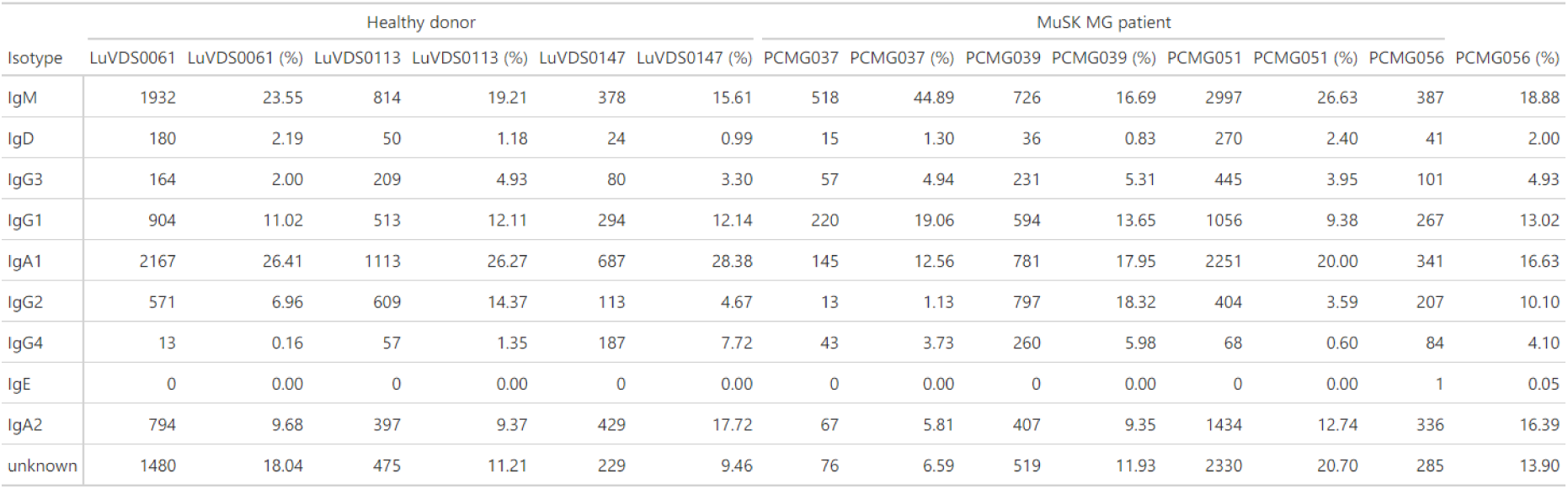
Isotype counts per cluster for each donor.

**Supplementary Table 6.** Isotype counts per cluster for each group.

| Isotype | Healthy donor |  | MuSK MG patient |  |
| --- | --- | --- | --- | --- |
|  | Count | Percentage | Count | Percentage |
| IgM | 3124 | 21.02 | 4628 | 24.60 |
| IgD | 254 | 1.71 | 362 | 1.92 |
| IgG3 | 453 | 3.05 | 834 | 4.43 |
| IgG1 | 1711 | 11.51 | 2137 | 11.36 |
| IgA1 | 3967 | 26.69 | 3518 | 18.70 |
| IgG2 | 1293 | 8.70 | 1421 | 7.55 |
| IgG4 | 257 | 1.73 | 455 | 2.42 |
| IgE | 0 | 0.00 | 1 | 0.01 |
| IgA2 | 1620 | 10.90 | 2244 | 11.93 |
| unknown | 2184 | 14.69 | 3210 | 17.07 |

**Supplementary Table 7.**
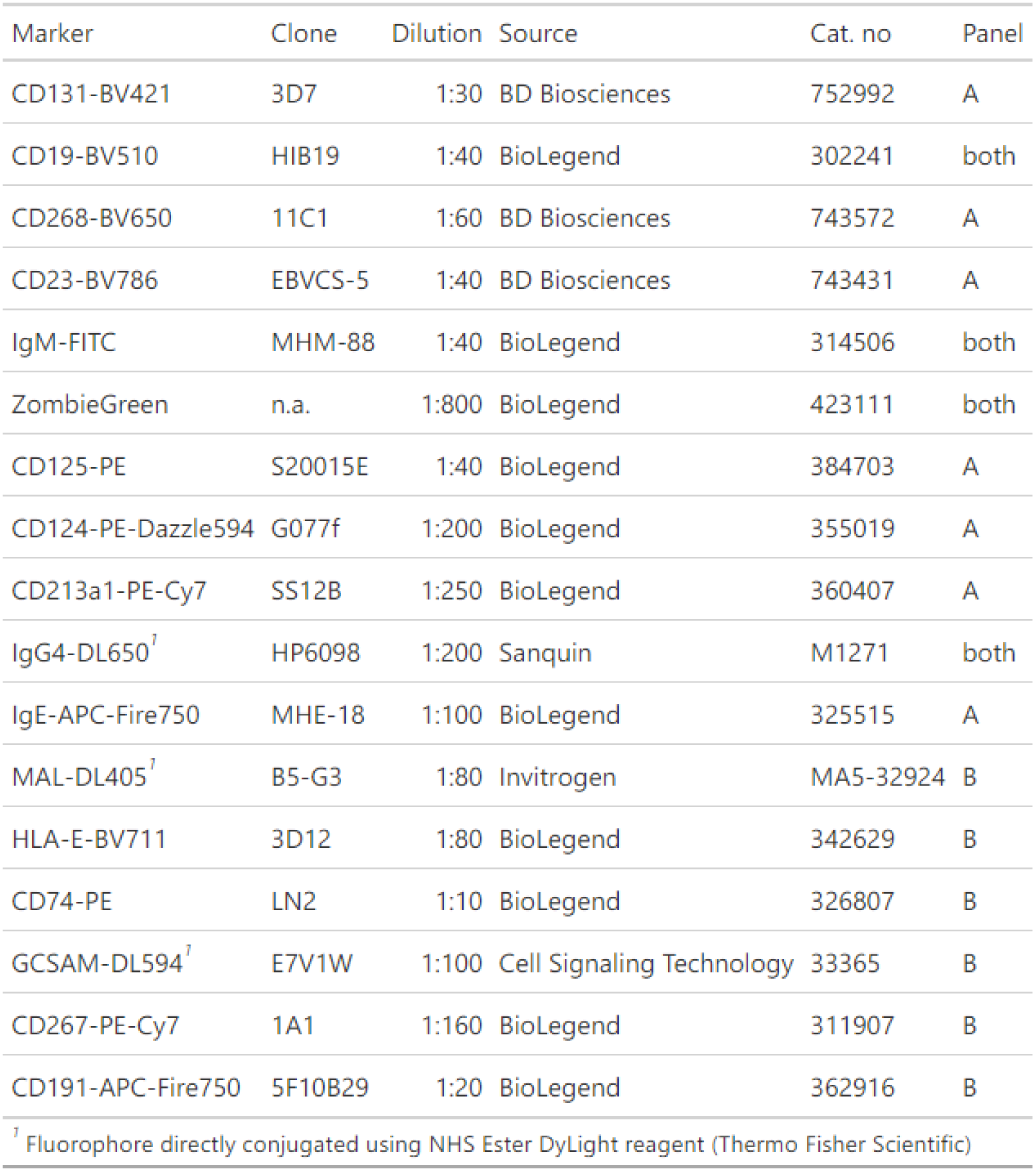
Flow cytometry antibody panels.

**Supplementary Table 8.**
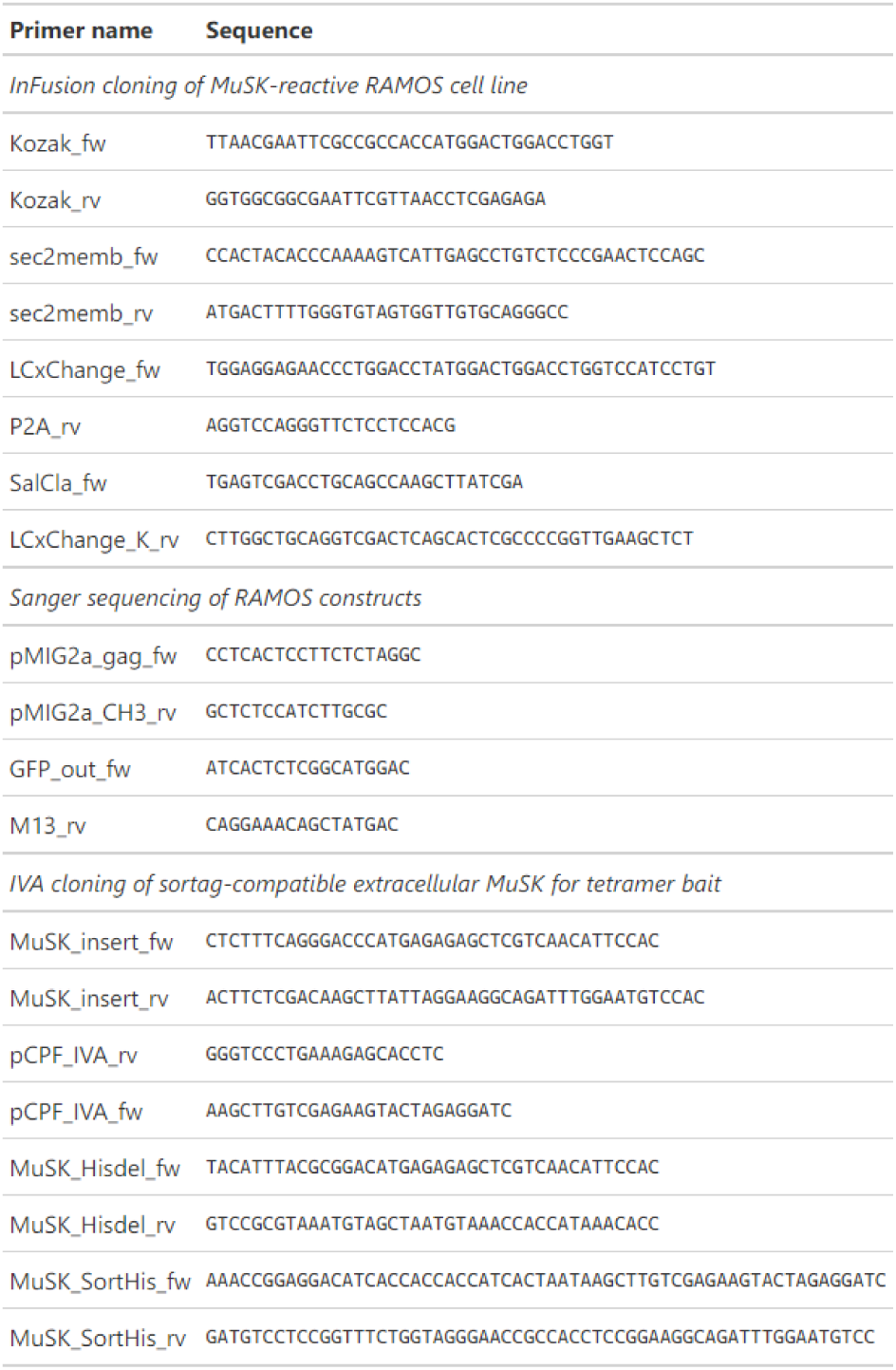
Primers.

**Supplementary Figure 1.**
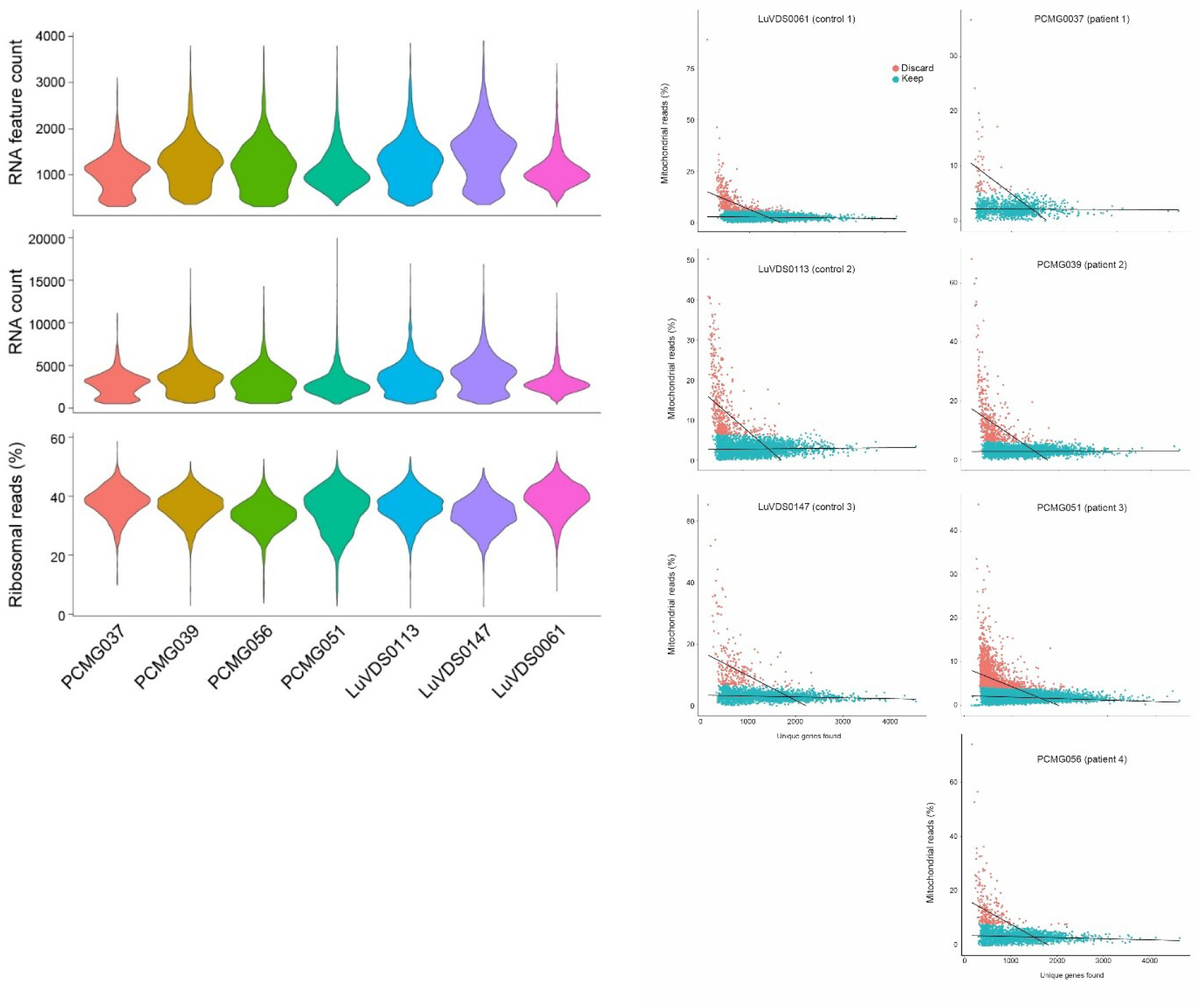
Single cell RNA seq quality control.

**Supplementary Figure 2.**
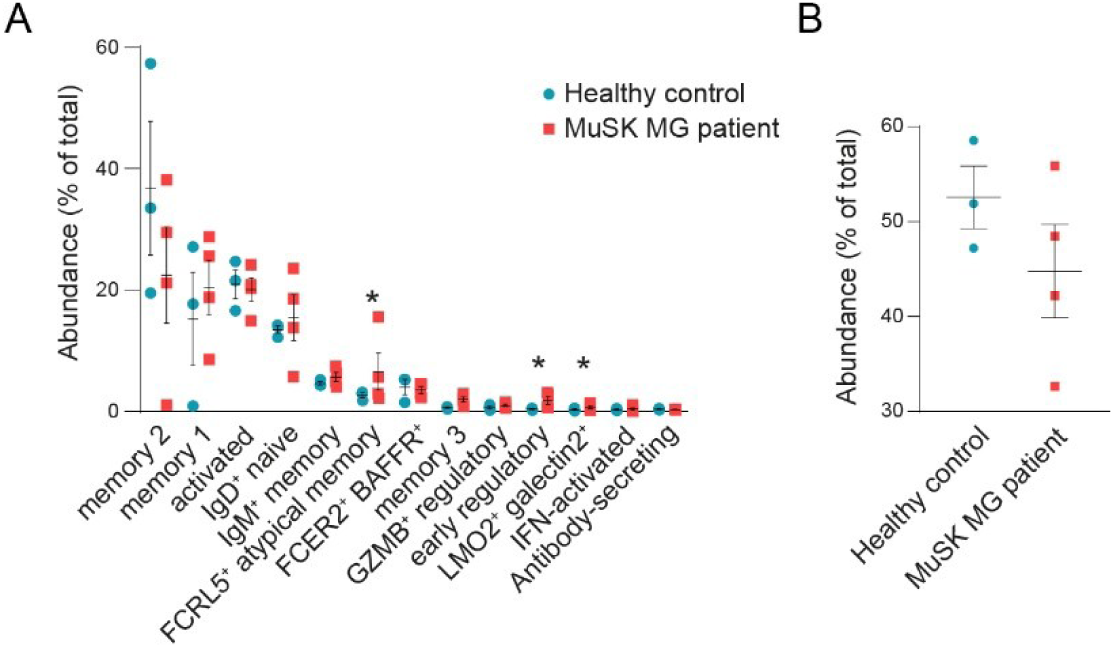
Relative abundance of MuSK MG patient and healthy control B cells for each cluster (A) and after merging 3 memory clusters (B). Multiple t-test with Šidák-Holm adjusted p-values (A), Student’s t-test (B). *: p < 0.05.

**Supplementary Figure 3.**
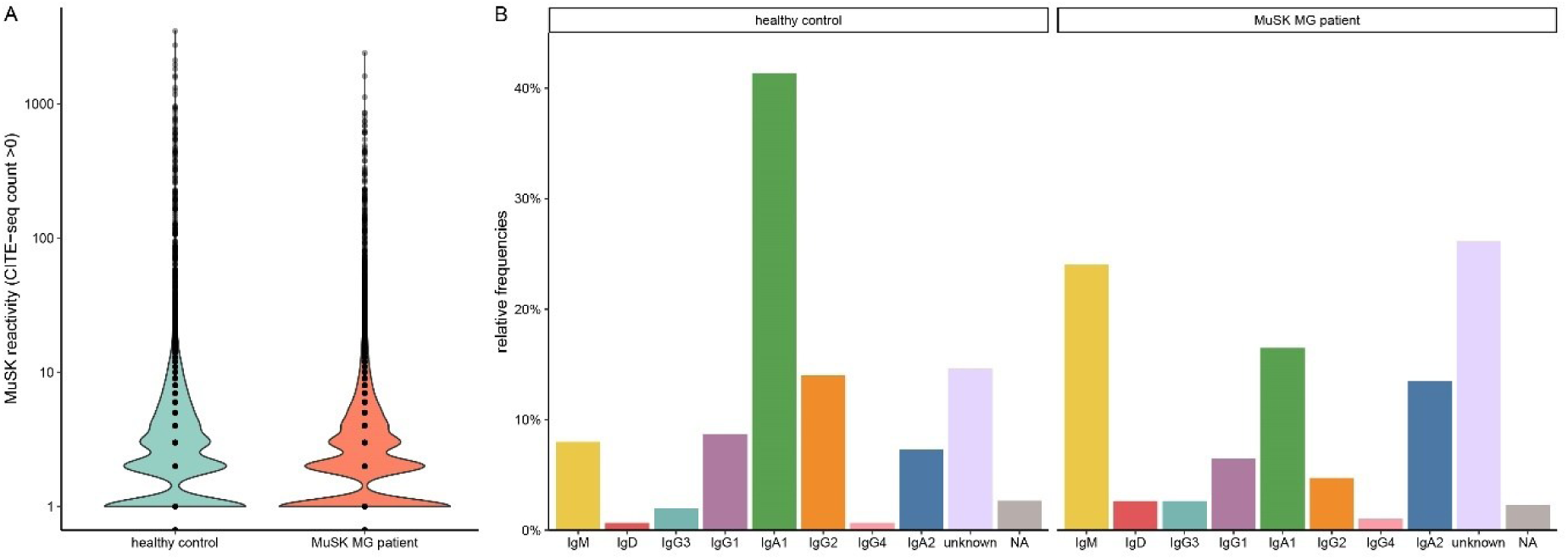
MuSK reactivity scores and isotype distribution in the GNLY^+^NKG7^+^ Bmem subsets.

**Supplementary Figure 4.**
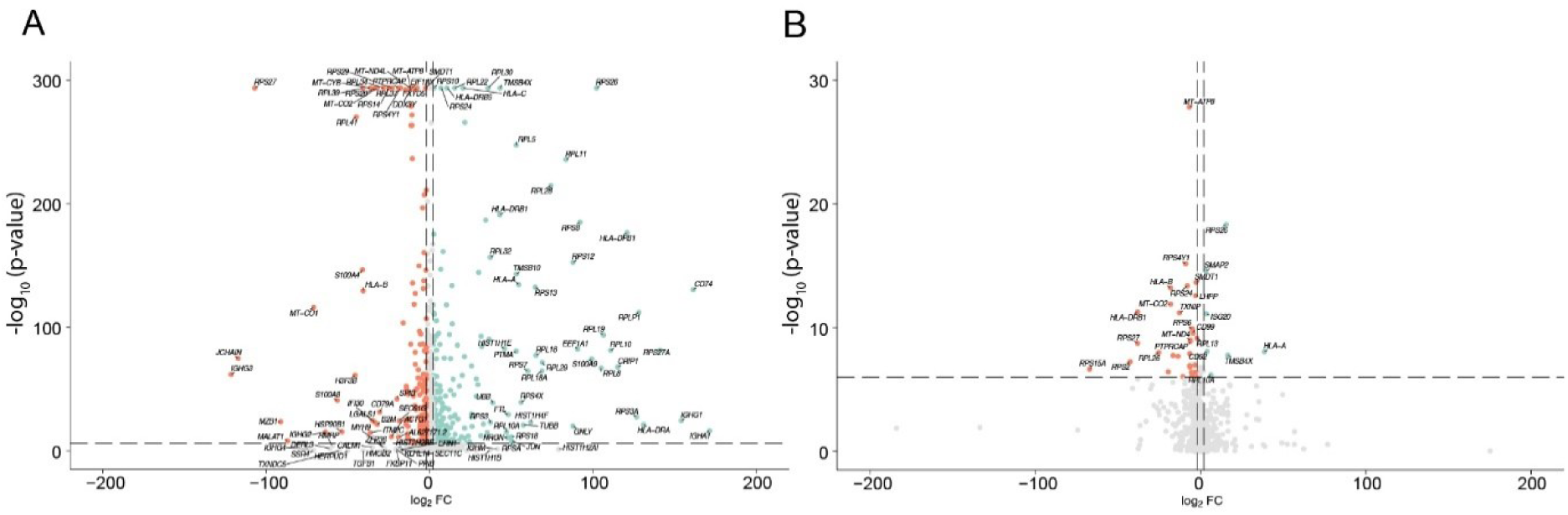
Differential gene expression of MuSK MG patient B cells over healthy controls in the total dataset (A) and specifically for IgG4^+^ Bmems (B).

**Supplementary Figure 5.**
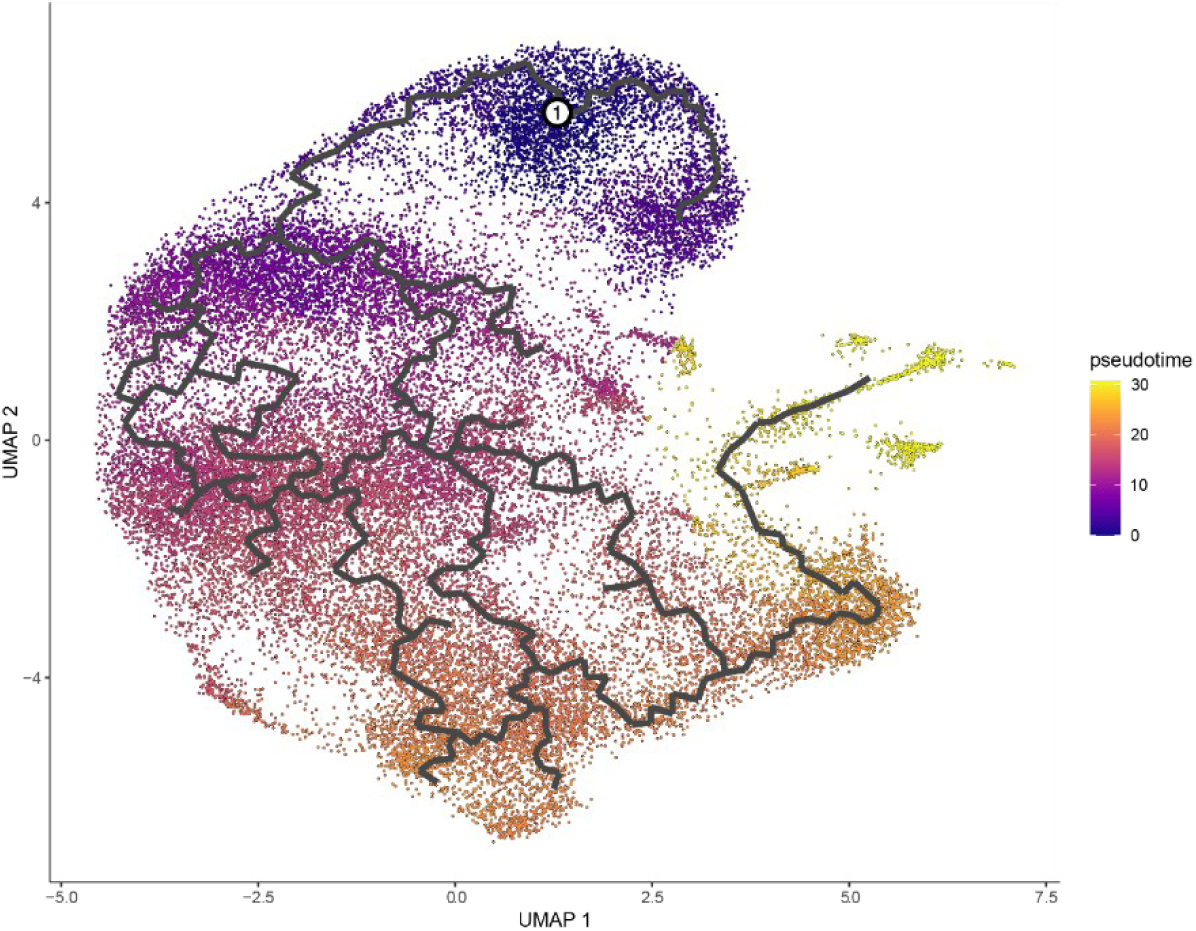
Pseudotime inference using *Monocle3* with IgD^+^ naive memory B cell cluster as starting node.

**Supplementary Figure 6.**
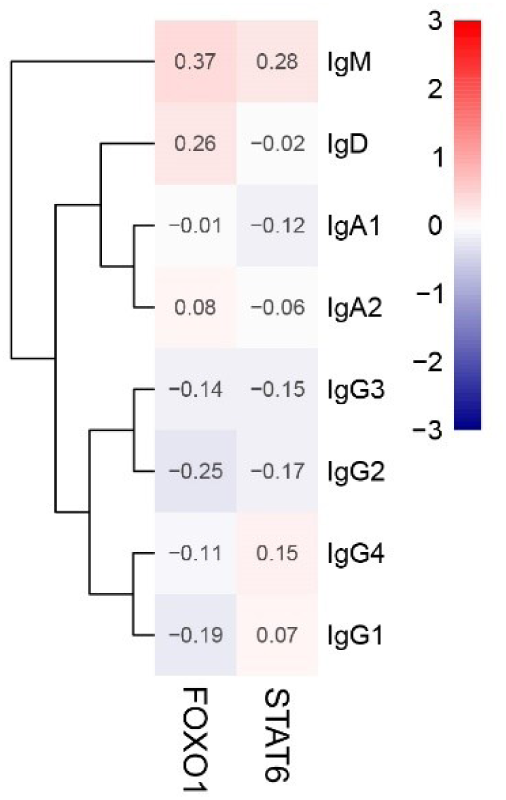
Inferred activity of transcription factors FOXO1 and STAT6 per isotype.

**Supplementary Figure 7.**
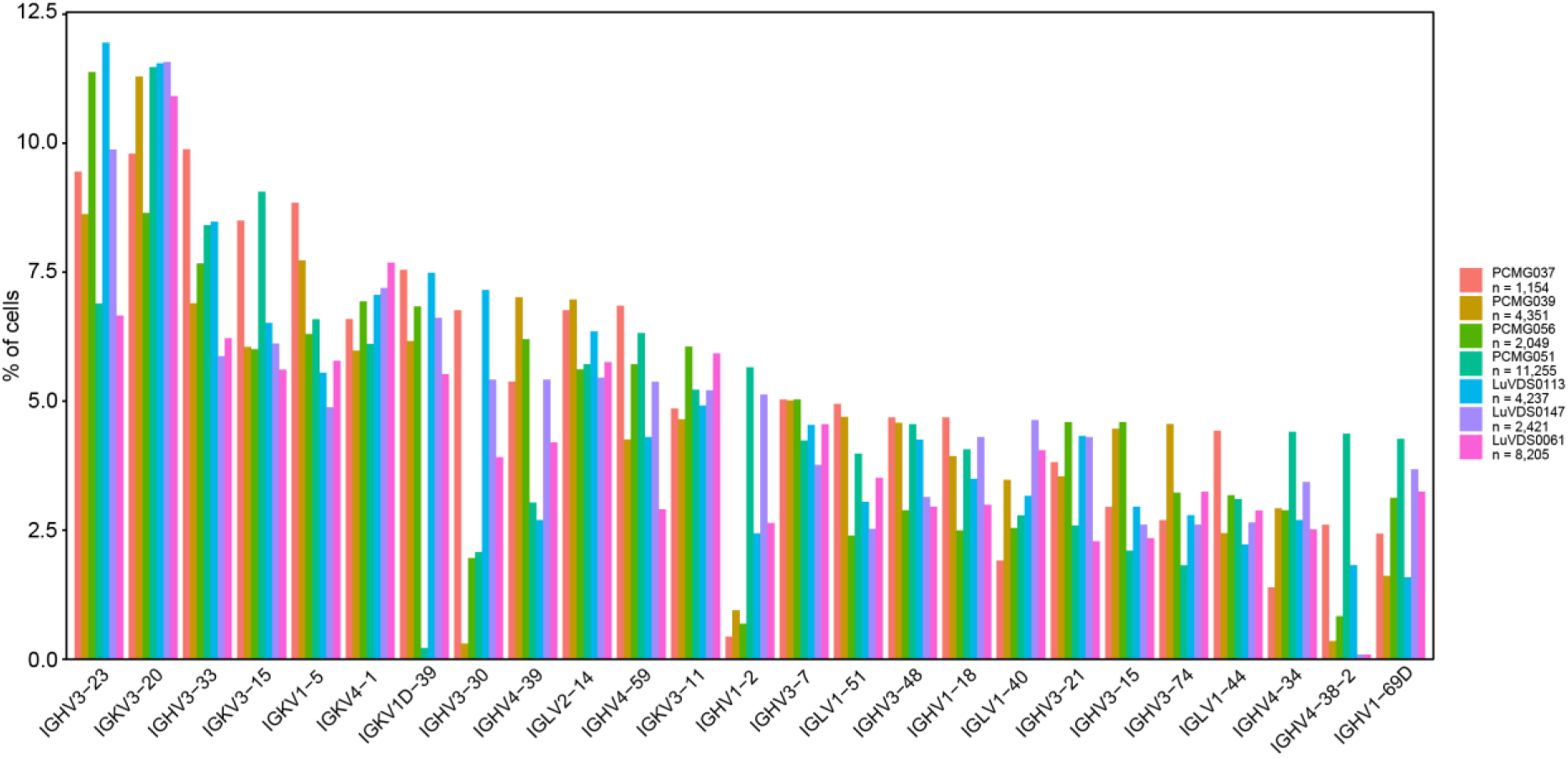
V-gene usage per donor.

**Supplementary Figure 8.**
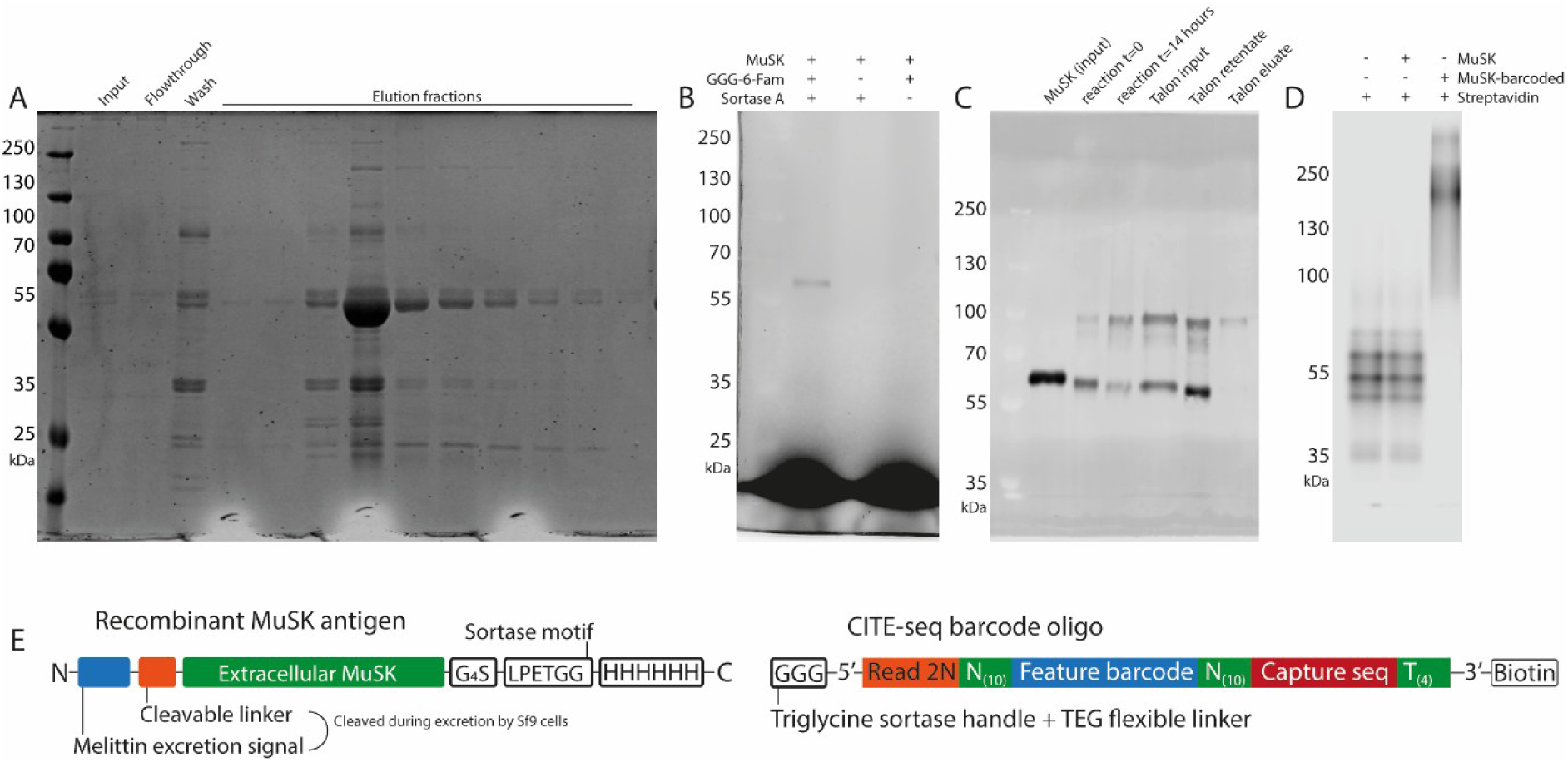
Recombinant MuSK tetrameric antigen. **(A)** FPLC purification fractions of recombinant MuSK from Sf9 cultures 72 hours post viral induction. Input: culture supernatant; Flowthrough: flowthrough over HisTrap FF Crude Ni column; Wash: column wash with 20 mM imidazole; Elution: 500 μl fractions of column elution with 200 mM imidazole. In-gel fluorescence image of SDS PAGE gel stained with PageBlue dye. Expected MW of recombinant MuSK antigen: 57.8 kDa. **(B)** Purified recombinant MuSK following 14 hour incubation with 6-fluoresceine amidite (FAM) modified with a triglycine repeat and Sortase A. In-gel fluorescence image of FAM fluorescence. Expected MW after sortagging reaction: 102.274 kDa (MuSK-6His: 57.8 kDa, DNA barcode: 45 kDa, biotin: 0.244 kDa, subtract GHHHHHH: 0.9 kDa). **(C)** Purified recombinant MuSK following incubation with triglycine repeat-modified DNA barcode and Sortase A pentamutant at t = 0 hours and t = 14 hours. Post-reaction cleanup by head-over-head incubation on Talon cobalt beads. Western blot image stained with mouse anti-rat MuSK primary antibody and IRDye800CW-conjugated donkeyanti-mouse secondary antibody. **(D)** CITE-seq barcode oligo-modified recombinant MuSK incubated with IRDye800CW-conjugated streptavidin. In-gel fluorescence image of IRDye800CW fluorescence. Expected size shift after complete MuSK-streptavidin tetramerization: 102.274 * 4 = 409.1 kDa. **(E)** Schematic representation of recombinant extracellular MuSK antigen and CITE-seq-compatible barcode oligo modified with 5’ triglycine repeat and 3’ biotin.

**Supplementary Figure 9.**
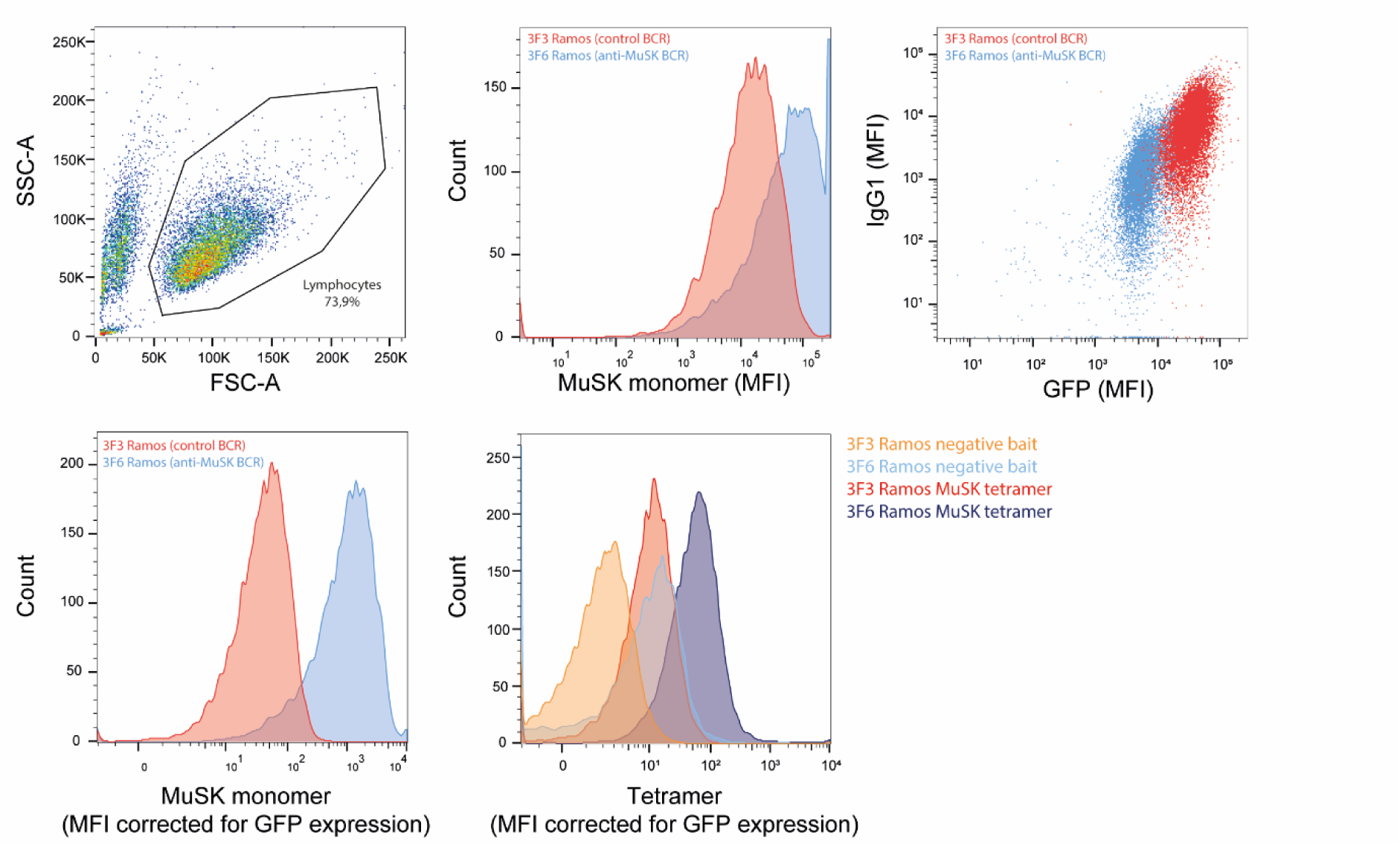
Validation of MuSK-reactive RAMOS cell line.

**Supplementary Figure 10.**
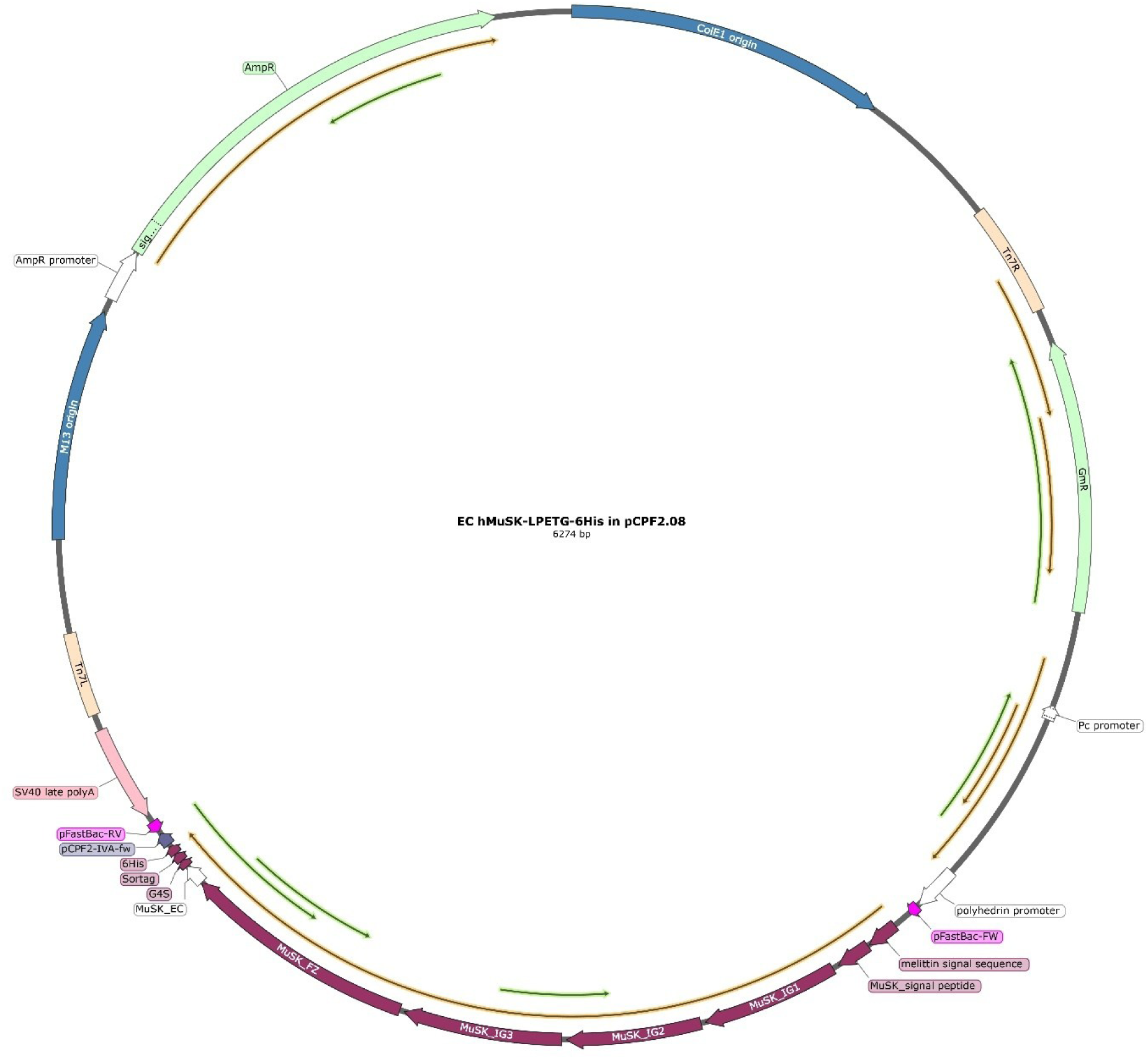
Plasmid map of recombinant extracellular sortaggable MuSK plasmid.

